# The ribosome directs nascent chains through two folding-dependent pathways

**DOI:** 10.1101/2025.04.08.647855

**Authors:** Alkistis N. Mitropoulou, T Tomasz Włodarski, Julian O. Streit, Sammy H.S. Chan, Eleni Plessa, Lauren F. Woodburn, Lisa D. Cabrita, John Christodoulou

**Affiliations:** Institute of Structural and Molecular Biology, University College London; Institute of Biochemistry and Biophysics, Polish Academy of Sciences, Warsaw, PL; Institute of Biochemistry and Biophysics, Birkbeck College, London, UK

**Author notes:** Correspondence: AM; LDC (l.cabrita@.ucl.ac.uk); JC. These authors contributed equally to this work (Alkistis N. Mitropoulou, T Tomasz Włodarski).

## Abstract

During their vectorial biosynthesis on the ribosome, elongating nascent polypeptide chains explore a range of conformational states towards their biologically functional structure. However, this high structural heterogeneity has limited their observation at high-resolution. Here, we have used an integrated structural biology approach to explore the structures of the multi-domain immunoglobulin-like FLN5-6 during its biosynthesis, capturing early folding through to native folding. We developed an in-silico purification approach for cryo-EM of ribosome-nascent chain complexes (RNCs), and integrated the resulting cryo-EM maps with NMR spectroscopy and atomistic molecular dynamics (MD) simulations to produce experimentally reweighted structural ensembles of RNC. The resulting atomistic structures reveal insights into the orientational heterogeneity of the nascent chain and its dynamic interactions with the ribosome. In particular, we find that two distinct pathways exist for nascent polypeptides in the exit tunnel vestibule, influenced by their stage of biosynthesis, folding conformational state and ribosomal RNA helices lining the tunnel. Our systematic analysis of the structures of nascent proteins translation-stalled at multiple time-points provides insights into how the ribosome dynamically modulates its pathway out of the exit tunnel to regulate its folding and accessibility for auxiliary factors of other co-translational events.

## Introduction

For the majority of proteins, folding is concurrent with their ribosomal translation^1^. While advances in protein structure predictions are increasingly providing exquisite descriptions of the equilibrium structures of biomolecules^2^, a molecular-level understanding of how structure, essential for biological activity and a healthy proteome, forms within cells remains obscure. Following peptide bond formation, nascent chains (NCs) navigate through the ribosome’s exit tunnel, while within the tunnel rudimentary structures can begin to form limited by spatial confinement^1,3,4^. Upon emergence from the exit vestibule, increasing freedom to sample conformational space exists^5–8^ and the dynamic process of co-translational folding (CoTF) ensues. Known to be influenced by several factors-NC interactions with the ribosomal surface^9,10^, steric exclusion arising from its proximity^3,9–11^, entropic forces exerted on the NC^12^, translational speed, as well as the engagement of molecular chaperones and other ribosome-associated factors^13–17^—the molecular details remain poorly understood.

When extended beyond the ribosome’s exit tunnel, solution NMR strategies have proved useful in characterising the structural dynamics of NCs^18–20^ while within the tunnel cryo-EM investigations remain the primary route to obtain structures: cryo-EM maps of RNCs have revealed detailed translational arrest mechanisms of stalling sequences^21–27^ and shown that α-helices can begin to fold in the ribosome exit tunnel^28^. More poorly defined NC electron densities have, however, been observed in visualisations of locally-structured segments of small protein tertiary structure motifs (< 100 aa) that can begin to fold within the ribosome’s upper exit tunnel and lower vestibule region^5,7,12,28–32^. Upon emergence from the ribosome exit tunnel, hydrogen-deuterium exchange mass spectrometry and NMR investigations have permitted complementary characterisation of the structure, dynamics, and energetics of RNCs^3,9,11,12,19,33–37^. Merging of experimental observations from both NMR^19^ and cryo-EM to restrain MD simulations has been particularly useful in delineating the NC’s interactions^30^ and conformational preferences^11,12^. The fifth domain of the tandem immunoglobulin-like filamin protein, FLN5, is used in these studies (and in the current work) and proved to be an invaluable model for understanding several unique aspects of coTF^3,12^ that are not observed in the folding of its isolated counterpart, including two obligate co-translational folding intermediate states (identified by ^19^F NMR^19^).

By applying a new cryo-EM in silico purification approach and 3D classification to FLN5-RNCs we capture here the earliest, intermediate and late stages of co-translational folding of FLN5. We deconvolute the NC’s extensive heterogeneity into a series of maps, dissecting each RNC state. Unlike prior maps that show an average of an RNC within one static snapshot^7,29,32,38^, this cryo-EM strategy enables capture of the NC’s range of dynamic structures, which yields up to thirteen classes per RNC length. Our structures unveil a unique bifurcation point which delineates the existence of two distinct pathways through the tunnel accessible to the elongating NC. By integrating an all-atom simulations approach to model FLN5’s folded conformation we show that the NC’s transit through these pathways alters during the course of elongation. This is modulated by the structure formed by the solvent-exposed, tethered NC, and a communication relay via a set of novel interaction sites within the ribosome. The data presented show that there is significant promise for extracting the atomistic details of the several and distinct conformational states being sampled by NCs during co-translational processes by merging the structural states from cryo-EM and NMR^19^ to restrain MD simulations.

## Results

### RNC design for structural studies of co-translational protein folding on the ribosome

We produced a set of purified biosynthetic snapshots from *E.coli*^1,39^ to report on FLN5’s co-translational folding pathway on the ribosome (Fig. 1a) by investigating FLN5*+L* RNCs with a linker *L* corresponding to a short stalling motif arrest-enhanced SecM (AE1)^19^ and increasing lengths of FLN6 (Fig. 1a). FLN5+31, FLN5+34, FLN5+47 RNCs permit the progressive emergence of FLN5 and report on its structure acquisition, including capturing increasing populations of its two folding intermediates^19^ and its completed native fold (*i.e.,* at FLN5+47, no U state of FLN5 is observed by NMR (Table in Fig. 1b). Two folding-incompetent variants of FLN5+31 A3A3 and FLN5+47 RNCs Y719E^9,15^ were also examined (Fig. 1a,b).

**Figure 1.**
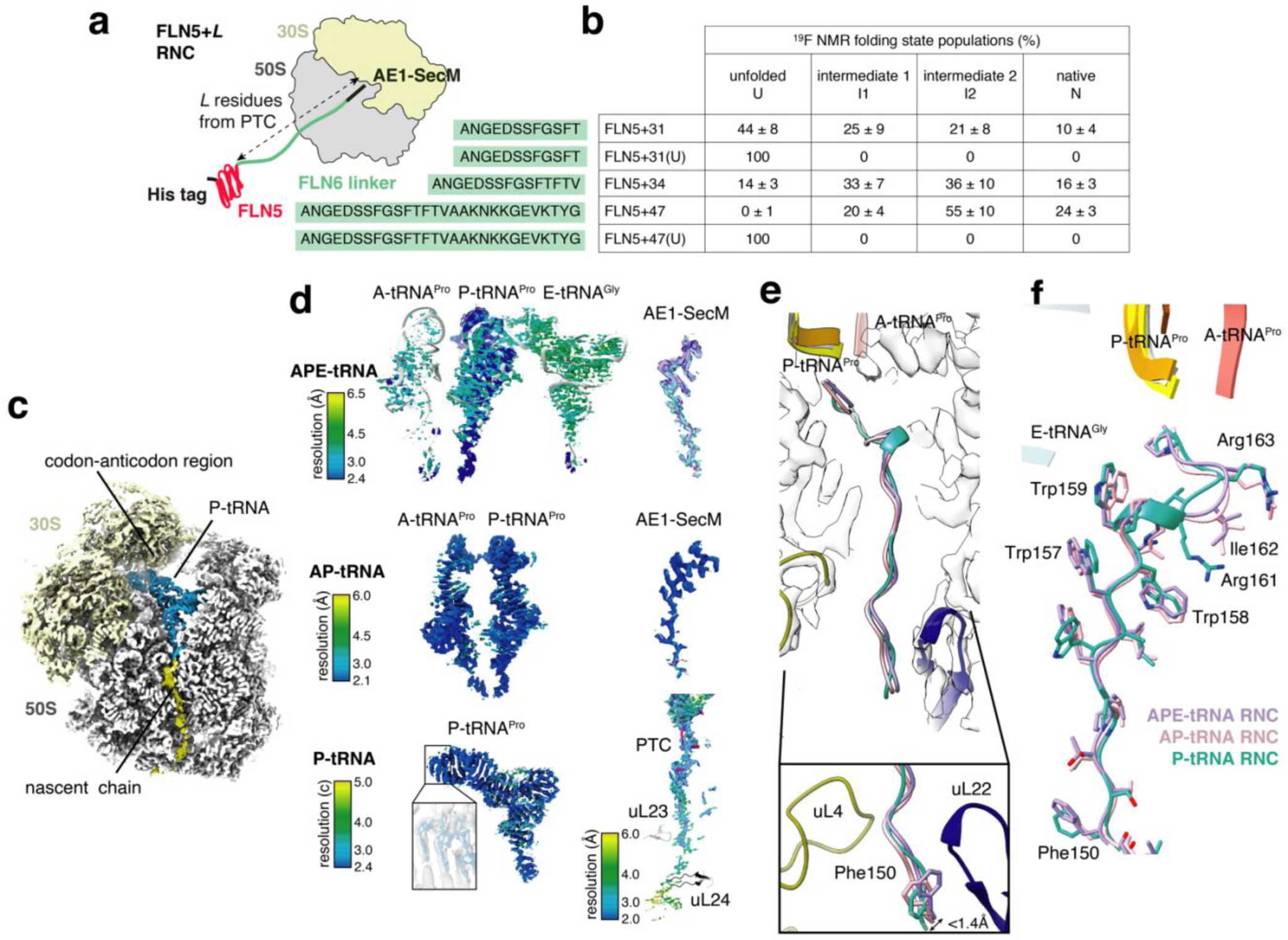
Cryo-electron microscopy maps of FLN5 biosynthetic snapshots capture three translocation states. **a**. Schematic designs of the SecM-arrested ribosome-nascent chain complexes of FLN5+L with increasing length L derived from the sequence of the subsequent FLN6 domain (in green) along with the C-terminal arrest-enhanced SecM stalling motif (FLN5+L) **b**. Table reporting on population of FLN5 states for the different linker lengths (L) as shown by 19F NMR data^19^ **c**. Density map of FLN5+34 RNC after the first 3D refinement showing 30S, 50S, P-tRNA and NC densities in beige, white, blue and yellow, respectively. **d**. The tRNA density for the three reconstructions of translocation states: APEtRNA (left), AP-tRNA (middle) and P-tRNA (right) RNCs coloured by local resolution. The tRNA structural models are shown in white cartoon representation. Density for the SecM-AE1 is shown for the APEtRNA and AP-tRNA RNCs, coloured by local resolution. An example of quality of codon-anticodon density obtained is shown for P-tRNA RNC alongside the RNC density coloured by local resolution, to illustrate the overall good quality of the maps. **e**. Superposition of all SecM-AE1 de novo built models for the FLN5+34 RNC from APEtRNA (purple), AP-tRNA (salmon) and P-tRNA (cyan) RNCs, presented within the ribosomal exit tunnel (white surface) spanning 50Å from the PTC and beyond the constriction site formed by uL4 (yellow) and uL22 (blue) loops. (inset) shows the position of the N-terminal amino acid of the SecM-AE1 sequence (Phe150) for all three RNCs. The pairwise RMSD is 0.4-1.3Å which is within the resolution limit >2A. **f**. Similar to c) in atoms representation (coloured by heteroatoms) to show the overall good alignment of SecM between the three translocation states.

### The nascent chain in the upper tunnel is virtually identical between the different translocation states

We focused our analysis initially on FLN5+34, as it is heterogeneously populated at equilibrium with unfolded and folded states, and partially-folded intermediate states in solution (Fig. 1b), although the FLN5 domain is entirely emerged from the exit tunnel^3^. *E.coli-*derived^39^, purified FLN5+34 RNCs were snap-frozen onto cryo-EM grids and subjected to data collection and processing^30^ (see Methods). An initial 3D reconstruction revealed the presence of unresolved, heterogenous tRNA cryo-EM density within the RNC maps, indicative of translating ribosomes having been captured in different translocation states (Fig. 1c and Supplementary Fig. 1a). A 3D classification with a mask in the inter-subunit space (See Methods, Supplementary Fig. 1a) identified three major classes with varying tRNA compositions and resulted in three distinct RNC reconstructions being obtained (Fig. 1d): (i) a pre-translocation state with only a P-site tRNA (Fig. 1d right) generated by 610K particles (P-tRNA RNCs); (ii) tRNAs present in both the A- and P-sites (pre-translocation state of A/A and P/P) from 280K particles (jointly referred to as AP-tRNA RNCs Fig. 1d middle & Supplementary Figure 2a); and (iii) tRNAs present in P- and E-sites with weak density in the A-site (300K particles), termed as APE-tRNA RNCs (Fig. 1d left, Supplementary Fig. 1a – for data processing pipeline). After CTF-refinement and 3D reconstruction in which the small ribosomal subunit (SSU) was subtracted, the initial average resolution of the ribosome (∼3Å) was further improved to average map resolutions of 2.1Å (P-tRNA RNC), 2.2Å (AP-tRNA RNC) and 2.5Å (APE-tRNA RNC) (Supplementary Fig. 1a, in purple, orange and cyan for APE-tRNA, AP-tRNA and P-tRNA RNCs). These high-quality maps obtained enabled visualisation of key peptidyl transferase centre (PTC) features, including discrete rRNA base-separation within the codon-anticodon interaction site (Fig. 1d right) and density for a proline attached to the A-tRNA (Supplementary Fig. 2b), as well as ten-nucleotide mRNA stretch near the tRNA anticodon (Supplementary Fig. 2a).

In all three reconstructions (P-tRNA, AP-tRNA and APE-tRNA RNCs) of FLN5+34 RNCs, the P-site tRNA was connected to continuous density corresponding to the sequence of the stalling motif SecM-AE1 (Phe150-Pro165) (sequence is shown in Supplementary Figure 2c) beginning at the PTC and traversing 50Å through to the uL4/uL22 constriction site (Fig.1c for the P-tRNA RNC and Supplementary Fig. 1b for the AP-tRNA (top) and APE-tRNA (bottom) RNCs). In all three maps, the C-terminal SecM-AE1 region (residues Trp157 to Pro165) near the PTC showed a higher local resolution (2.1-2.5Å) relative to its N-terminus (Phe150 to Ile156), located closer to the constriction site (2.5-3Å) (Fig. 1d). A side-chain model corresponding to the SecM-AE1 density was subsequently built *de novo* and apart from a small helical turn formed between Trp159 and Arg161 that is uniquely present in P-tRNA RNCs (Fig. 1e), SecM-AE1 is otherwise in an extended conformation and is unchanged across all tRNA translocation states (Cα RMSD values of 0.97Å (between P- and AP-tRNA RNC) and 0.95Å (between AP-tRNA and APE-tRNA RNCs)) (Fig. 1f). This observation includes the downward register of Phe150, the most N-terminal SecM-AE1 residue, whose position also remains unchanged in the tunnel (Cα RMSD values of 0.4 and 1.3Å in AP-tRNA and APE-tRNA versus P-tRNA as reference, respectively) within the resolution limits (<2.5Å) of the maps (Fig. 1e).

### *In silico* purification reveals a range of NC conformations within the exit tunnel

Within the middle tunnel region, between 50-80Å from the PTC (Fig. 2a), enclosed by the constriction site and ribosomal protein uL23, continuous NC density attributable to FLN6 is observed across all translocation states of FLN5+34 RNC (P-tRNA RNC is shown as an example in Fig. 2a & Supplementary Fig. 1b for the AP-tRNA (top) and APE-tRNA (bottom) RNCs). The local resolution of the NC in the tunnel varies considerably (from 2Å at the narrowest point to 7Å at the widest, Fig. 2b), the latter consistent with the increased dynamic conformational freedom within this widened region (radius ∼6-8 Å) relative to the constriction site (radius 5Å)^40^. Beyond this, in the wider vestibule region (80-100 Å from the PTC) which is located between the protruding loops of uL23 and uL24, the NC density becomes substantially sparser and is discontinuous (at 1.5σ level). Moreover, no NC density can be traced between helix 6 (H6) and H7 of 23S rRNA along one side of the vestibule and helices H24 and H50 on the opposing face of the tunnel although some NC observability is regained further down, close to H24 of 23S rRNA (Fig. 2a & Supplementary Fig. 1b). NC density next becomes traceable at the very end of the vestibule at the tip of uL24 loop (which consists of residues Pro48-Pro55).

**Figure 2.**
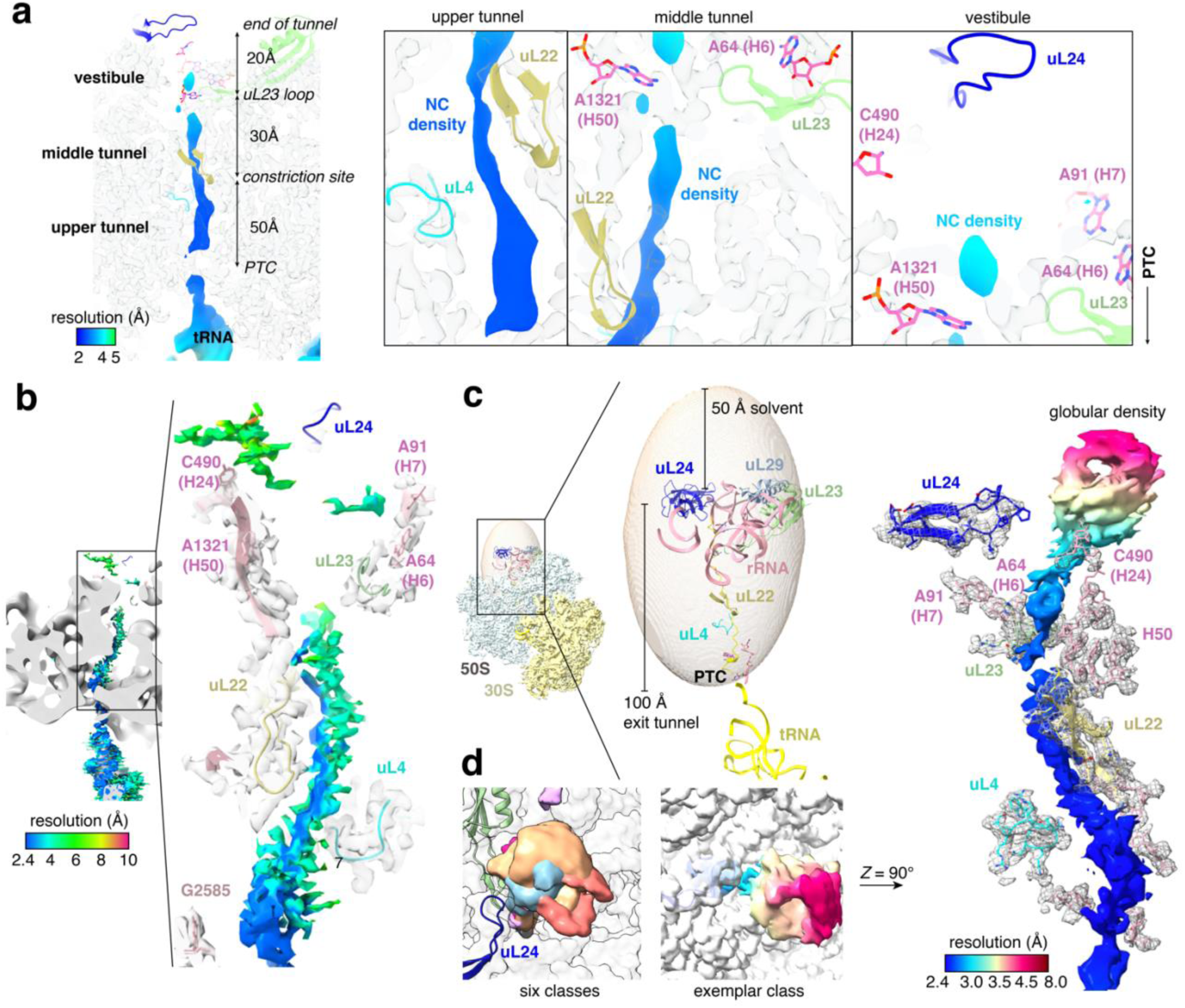
FLN5+34 RNC reveals different NC states with globular domain density outside the tunnel and an NC path bifurcation point within the exit tunnel*. a.* Cross-section of the 50S subunit of FLN5+34 RNC reconstruction coloured according to local resolution. The ribosome density is shown in white and important tunnel components are highlighted in pink atom representation for rRNA bases and cartoon for uL4 (cyan), uL22 (yellow), uL23 (green) and uL24(blue). through the exit tunnel. The three main regions of the tunnel are indicated and their magnified versions are shown on the right. **b**. FLN5+34 NC coloured according to local resolution before in silico purification. Landmarks for the ribosomal tunnel wall are also shown: 23S rRNA (pink), uL4 (cyan), uL22 (yellow), uL23 (light green) and uL24 (blue). (inset, right) Magnified view of the density corresponding to the NC from the PTC through to the vestibule and beyond where density is sparse. Ribosomal landmarks are also shown for guidance. **c**. For tunnel 3D classification of the RNC particles an ellipsoid mask was created that included the tunnel and are surrounding the tunnel from the PTC to the end of the tunnel and 50Å beyond. The 50S and 30S density is shown in light blue and yellow, rRNA is shown in pink and ribosomal proteins are in cartoon. Right: zoom in the ellipsoid mask and ribosomal tunnel components. **d**. A superposition of twelve NC models within the ribosomal exit tunnel, as built from cryo-EM density. Ribosomal landmarks shown include uL4 (cyan), uL22 (beige), uL23 (light green) and uL24 (blue), 23S rRNA bases A64-A90 (H6-H7 rRNA helices), C490 (H24 rRNA helix) and A1321 (H50 rRNA helix) shown in pink atoms. Middle panel: 90° rotation view looking through the tunnel showing one representative class coloured according to local resolution. Right panel: zoom into the NC density showing improvement after 3D classification compared to b.

Beyond the tunnel exit and close to the surface of the ribosome, sparse NC density (>8Å resolution, σ=1.5) is observable (Supplementary Fig. 1b) across all translocation states of FLN5+34 RNCs. Upon closer inspection, this globular density is observed in the vicinity of the vestibule and is consistent with the presence of either a natively- or partially-folded FLN5 domain. To further deconvolute the NC density in FLN5+34 RNCs, the maps were subjected to a new *in silico* purification method, which involves a 3D classification of the exit tunnel (flow-scheme shown in Supplementary Fig. 1a, and Methods). For this *in silico* purification, we developed a mask whose shape enveloped the entire exit tunnel and extended *ca.* 50Å beyond (Fig.2c) the ribosomal exit (the latter boundary as defined by the uL24 loop). The mask included a substantial portion of the 23S rRNA core (defined by a radius of 17Å from the centre of the tunnel and was identical between reconstructions) that encases the tunnel. Then, following a round of focussed 3D classification, twelve subclasses, six for each of the P-tRNA (class-1 to 6) and AP-tRNA (class-7 to 12), were obtained for FLN5+34 RNCs (Fig. 2d for the models, Supplementary Fig. 1a shows the classification process and Supplementary Fig. 3 shows densities for each of the classes); these indicate average resolutions of 2.2-2.8Å for all the classes (from 26K to 171K particles). The improved resolution of these maps allowed continuous NC density to be visible throughout the entire length of the tunnel and beyond the exit (Fig. 2b middle and right density and Supplementary Fig. 3). *De novo* model building of FLN6 residues showed extended conformations of this segment of the NC spanning from uL22, after the constriction site, to the end of the vestibule near uL24, but superposition of the FLN6 models across all maps showed variability in their conformations (Fig. 2d), averaging 5Å Cα RMSD from Ala751 to Val765 (*c.f.,* SecM-AE1 Cα RMSD 0.4-1.3Å). Evidently, the wider region (∼8Å radius) of the tunnel vestibule allows the NC to explore a far greater range of conformations relative to the narrower confines (4-6Å radius) of the upper tunnel near the PTC. Beyond the exit, the globular-like density (where the FLN5 domain is expected to be located) could not be entirely satisfied with *de novo* model building alone.

### All-atom molecular dynamics simulations complement structural analyses of cryo-EM maps

To overcome the challenge of interpreting the RNC globular density beyond the tunnel, we performed unbiased, all-atom MD simulations of folded RNCs of FLN5+34 (as well as FLN5+31 and FLN5+47 that are discussed further below) with explicit solvent (8 μs total simulation time for each construct, see Methods). Since the globular density can accommodate folded FLN5 and models of the intermediate states are not available we modelled only the native state. These simulations sample a range of FLN5 orientations and ribosome interactions and show that FLN5-ribosome interactions decrease with increasing linker length (Supplementary Fig. 4) as expected based on NMR measurements (Supplementary Fig. 5). We reasoned that the cryo-EM maps represent local minima on the FLN5 RNC energy landscape, which we aimed to sample in the simulations. Thus, to improve agreement between MD-derived structural states and NC maps, we applied cryoENsemble^41^, an iterative Bayesian-reweighting methodology (Methods & Supplementary Figure 6). In all cases, we find that the reweighted sub-ensembles better represent the cryo-EM maps than a single, best structure or the initial ensemble (Supplementary Fig. 7). This suggests that the simulations have successfully captured the relevant FLN5 orientations and ribosome binding sites present in the cryo-EM data, and that these data may be described by native or at least near-native FLN5 conformations. The resulting structural models are discussed in the subsequent paragraphs.

### The nascent chain bifurcates in the tunnel via two exit pathways modulated by rRNA

Across the 12 cryo-EM RNC classes of FLN5+34 described above, the vestibular NC assumes several distinct conformations (Fig. 2d & Supplementary Fig. 8): within the lower tunnel, FLN6 residues Ala751-Val765 are primarily observed along one face of the tunnel, delineable between rRNA H24 and H50, and the uL24 loop (Fig. 2d). Along this pathway, the NC makes contacts with the tunnel including a distinct interaction with nucleotide C490 (Fig. 3a & Supplementary Fig. 9) of the H24 helix, and with other rRNA in the vicinity of this area (as observed in Classes 1, 5, 6, 7, 9 and 11). Contacts are also observed for the majority of classes between H50 (A1321) and Thr764 and Phe761 of FLN5-6.

**Figure 3.**
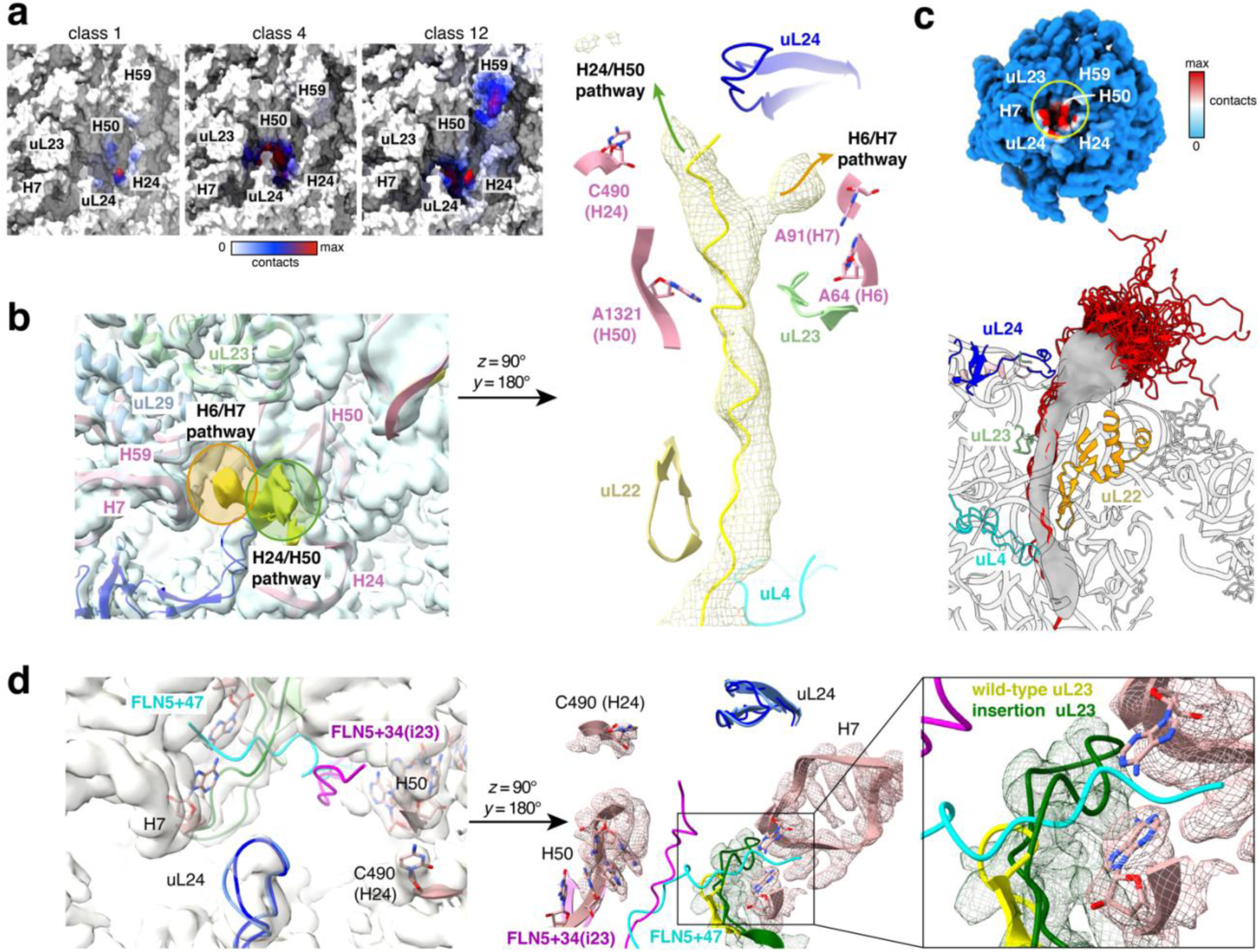
The NC follows two distinct pathways in the vestibule. **a**. Contacts between the ribosomal vestibule and the NC density. White colour (no contacts) means no NC density voxels were observed within 10Å from this part of the ribosome vestibule and red (Max contacts) colour indicates that this part of the ribosome surface has the highest number of NC density voxels observed near it. **b**. Left: Two pathways observed for density in vestibule in the P-tRNA_4 FLN5+34 RNC class. The NC cryo-EM density (shown in yellow upon Gaussian filtering (σ =1) bifurcates beyond the uL23 with one pathway near H6/H7 free base pair and a second pathway near H50 and H24. uL23(light green), uL24(blue), uL29 (light blue) and specified 23S rRNA bases (pink) are highlighted. A surface representation of the 50S is depicted in transparent light blue. Middle: shows P-tRNA_4 FLN5_34 RNC as rotated 90 degrees around the z-axis. The cryo-EM density difference map (yellow mesh) shows the two pathways, and the most prevalent NC pathway is depicted (yellow cartoon). Right: schematic depicting the location of the two pathways. **c**. Reweighted all-atom molecular dynamics simulations capture FLN5 structural heterogeneity. Left: the MD-derived FLN5-6 NC models selected upon cryoENsemble reweighting with cryo-EM maps (class-7 from +34) are shown in red cartoon representation along with the nascent chain cryo-EM maps (Gaussian filtered) in grey and ribosome cross-section (with uL4, uL22, uL23, uL24 and uL29 proteins highlighted in cyan, orange, light green, blue and pink, respectively). Analysed ribosome-NC binding interactions were mapped on the ribosome surface (right). **d**. CRISPR-modified uL23 loop (uL23 insertion is shown in dark grey and wild-type uL23 is shown in yellow) blocks the NC pathway via rRNA H6/H7. The FLN5+47 class 2 NC (cyan) has been included to show the NC path via H6/H7 compared to the FLN5+34i23 NC (in magenta) which exits via H24/H50 path. Left: ribosome view from the top (with uL23, uL24 and rRNA highlighted in green (and yellow for the wild-type 70S), blue (and light blue for the wild-type 70S) and pink, respectively). Right: z-rotated view of the left image and inlet shows a zoom in version of the uL23 loop with an NC that would clash in the presence of the inserted loop. The RNC density is only shown for the rRNA and uL23 for clarity

Alongside this ‘H24/H50 pathway’, the highly populated class-4 reconstruction (Fig. 3b) (∼20% of all particles, and at 2.1Å, having the highest average resolution) shows additional FLN6 density consistent within a second discernible pathway (Fig. 3b). This pathway lies on the opposite face of the tunnel relative to H24/H50 and traces along the H6/H7 helices of 23S rRNA and in close proximity to uL23 (Fig. 3b). The observation of these two NC pathways, with the NC either along H24/H50 or H6/H7 at the exit port of the vestibule suggests a distinct bifurcation point in the tunnel. Further inspection traces the origin of this bifurcation to the 8-residue uL23 protruding loop (Fig. 3b middle and right panels). Attempts to further deconvolute the two NC pathways in class-4 (via Relion^42^, CryoSPARC 3DVar^43^ and CryoDRGN^44^) proved not possible, actually suggesting the co-existence of just the two distinct, parallel NC pathways. Of the two pathways, H24/H50 is apparently preferred for FLN5+34 since the FLN6 NC density is present in all classes and remains visible even at high threshold levels of the map (> 5σ); this is in contrast to the H6/H7 pathway which is only observed at lower density threshold (2σ) and is present in only one class (Supplementary Fig. 3 class 4).

The region corresponding to the two putative pathways were also independently observed in the FLN5+34 RNC MD simulations before reweighting (Supplementary Fig 10). Upon subsequent reweighting with cryoENsemble, the NC structural subensembles exhibit improved agreement with the globular NC density beyond the tunnel (Fig. 3c left & Supplementary Fig. 11). We can observe FLN5 domain interactions with the ribosome surface localised around the uL24 loop and with rRNA helices H24, H50 and H59 (Fig. 3c right). The folded domain is also interacting with H7 and uL29/uL23, but the NC linker is not observed along the H6/H7 pathway (Supplementary Fig. 12).

### Disruption of NC interactions with the A61-A93 base pair of rRNA can shift the pathway of the nascent chain

These combined observations hinted at a role of specific regions of rRNA in modulating the NC’s transit through the tunnel. To investigate this further, we addressed the question of how an NC’s course inside the tunnel changes when one of the pathways is obstructed. We thus compared the cryo-EM structures of FLN5+34 RNC (described above) with that of an engineered ribosome variant, 23i, which harbours an 8-amino acid extension to the uL23 loop that shields the H6/H7 regions and promotes earlier native folding of FLN5 RNCs (FLN5+34_23i RNCs)^30^.

On applying the *in silico* tunnel 3D-classification to cryo-EM data of FLN5+34_23i RNCs^30^ the NC density is observed solely along the H24/H50 pathway (Fig. 3d). A comparison of the entire tunnel-lining ribosomal components in the two structures (FLN5+34 and FLN5+34_23i) reveals them to be virtually identical (RMSD < 0.8Å, see Methods) with the important exception of observations in the conformations of A64 (H6) and A91 (H7) in FLN5+34_23iRNCs, which showed RMSDs of 1.2Å and 3.3Å respectively (Supplementary Fig. 8). Both H6 and H7 interact with the extended uL23 loop through its increased net positive charge (pI 8.3, vs. 7.3 in wild-type ribosomes), and the extension obscures the H6/H7 contact sites of the NC (Fig. 3d zoom & Supplementary Fig. 7). This apparent occlusion of the H6/H7 pathway in FLN5+34_23i may be a factor in inducing the observed earlier onset of NC folding in this variant^30^. The FLN5+34_23i RNC structure shows that this appears to be achieved by altering the path flux of the NC towards the H24/H50 pathway, which is favoured by folding-competent NC states at this stage of biosynthesis.

### The dominant NC pathway in the vestibule switches during biosynthesis according to its folding status

We next determined the cryo-EM structure of FLN5+47 RNCs (schematic in Fig. 1a). This species represents a subsequent snapshot in the biosynthesis of FLN5-6, to further investigate the role of flux within the tunnel in how a NC adopts a particular pathway. In solution, FLN5+47 RNCs show no observable unfolded FLN5 state and instead populate a range of structured states including native (∼25%) and partially-folded intermediate states (I1 ∼20% and I2 ∼55%)^19^. Following *in silico* purification of the cryo-EM FLN5+47 RNC data, nine classes could be resolved (Supplementary Fig. 13a,b). The reconstructions revealed NC density in the tunnel with a resolution ranging from 2.9Å (SecM) to 6-7.5Å (towards the uL23 loop) (Supplementary Figure 13c). In all classes of FLN5+47 RNC, the resulting *de novo* models are seen to exclusively follow only the H6/H7 pathway (Fig.4a).

The three C-terminal FLN6 residues (Ile776-Gly778) were found to form an α-helical turn in all RNCs (Fig. 4a), just after the constriction site formed by uL4 and uL22 loops. Residues Lys770-Phe761 are seen to interact with the uL23 loop and A91 from rRNA H7 (Fig. 4b for indicative examples) where hydrogen bonds and hydrophobic contacts can be observed, respectively. Pairwise comparison of the Cα atom positions of *de-novo* built FLN6 (residues Phe763-Gly778) shows an average RMSD of 4Å (RMSD of residues A751-V765 of all FLN5+34 RNCs is 5Å), among the classes indicating no discernible differences between the FLN6 pathways (of observable residues) in the vestibule within the resolution limits.

**Figure 4.**
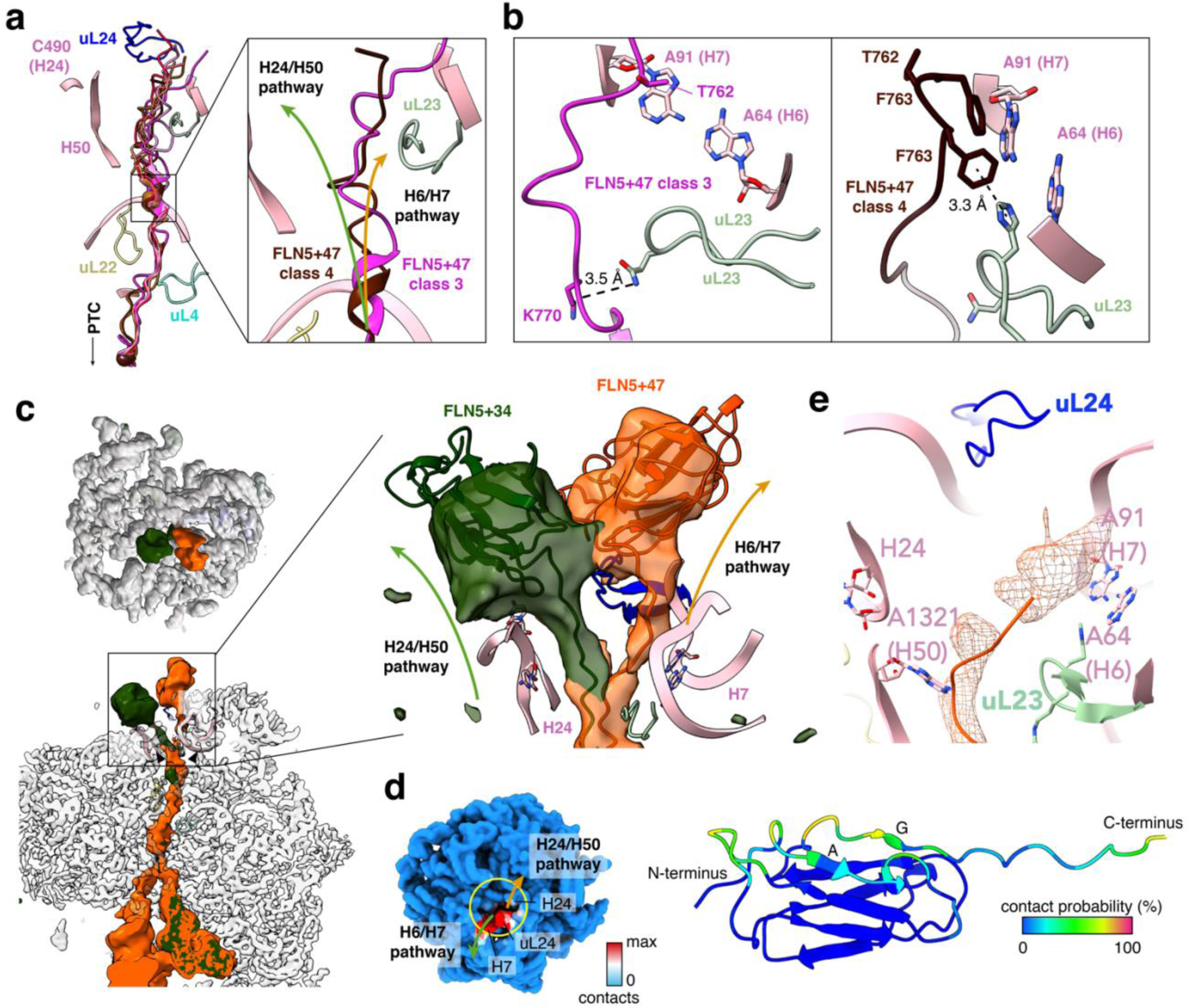
The NC exit path is related to the presence of globular density outside the tunnel. **a.** The FLN5+47 nine NC models within the ribosomal exit tunnel as shown with the following ribosomal landmarks; uL4 (cyan), uL22 (beige), uL23 (light green) and uL24 (blue), 23S rRNA bases A64-A90 (H6-H7 rRNA helices), C490 (H24 rRNA helix) and A1321 (H50 rRNA helix) shown in pink atoms. Right: zoom in the area of the middle tunnel and part of vestibule showing representative classes-3 and −4 depicted in magenta and brown, respectively. **b.** Interactions between class-3 (magenta) and class-4 (brown) with uL23 and rRNA helices H6/H7. Hydrogen bonds are shown in black dashed lines and π-π stacking is shown in black line. Residues involved in interactions are shown in atoms and coloured by heteroatom. **c**. Left: the NC cryo-EM densities of FLN5+47 and FLN5+34 RNCs are shown in orange and green, respectively. Middle: an overlay of FLN5+34 RNC (green) and FLN5+47 RNCs (orange) showing the comparison between two RNCs cryo-EM densities. Note that two RNCs follow different pathways inside the tunnel. Right: zoom into the vestibule region showing the potential bifurcation point between H50 and the uL23 loop. the two different length RNCs followed virtually identical pathways from PTC until the bifurcation point. The models for FLN5+34 and FLN5+47 were manually built for AE1-SecM and FLN6 (where density quality permitted) whereas FLN5 domain (from X-ray structure^58^) was rigid-body fitted and refined with Isolde or jelly-body fitted for the FLN5+34 or FLN5+47, respectively. Both models depicted here for illustrative purposes only. The globular domain densities will be explained later with ensembles as they represent more than one models each. **d.** Reweighted all-atom molecular dynamics simulations for FLN5+47; analysed ribosome-NC binding interactions were mapped on the ribosome surface (left) and on the FLN5-6 structure (right). **e**. Shows the cryo-EM density of the unfolded FLN5 RNC variant FLN5+47^U^ (in orange) (at 1.5 σ). The NC interacts with H6-H7 nucleotides; no density is observed further in the vestibule or beyond. Right: zoom into the vestibule area, showing the polyalanine model of FLN5+47^U^ in red cartoon and the NC cryo-EM density at 1.5σ

A comparison of the cryo-EM maps of these two snapshots (FLN5+34 and FLN5+47) shows that the NC appears to be able to follow two distinct pathways within the vestibule (Fig. 4c). At the earlier snapshot (L=34), that in solution populates a small unfolded population (14%) amongst the majority population (> 85%) of structured states (N, I1, I2), the NC follows the H24/H50 pathway in at least 80% of the particles. In these RNCs the NC is seen to be compact i.e. at least partially folded and partially inside the vestibule close to the uL24 loop (Fig. 2d middle panel).

The vestibule shape accessible for the NC when it follows the H24/H50 path (Fig. 3c) would appear to be sufficient for accommodation of such structure (given the short linker length, L=34) within the tunnel. FLN5+47-RNCs harbour an additional 13 FLN6 residues and show FLN5 globular density to be located only well outside of the vestibule in a range of positions and orientations (Supplementary Fig. 11b shows the location of globular domain outside the tunnel) within the area above the tip of the uL24 loop and rRNA helices H50, H59 and H24. In these RNCs, the FLN6 residues (Ser760-Gly778) are observed along the H6/H7 pathway within the tunnel (Supplementary Fig. 11b), while others (those closer to the N-terminus of FLN6, Ala751-Gly759) show only sparse density. The FLN5-attributable globular density is beyond the vestibule and not constrained by the spatial restriction of the tunnel. CryoENsemble reweighting of the FLN5+34 and FLN5+47 MD ensembles using these cryo-EM maps provide further insights; the FLN5+47 NC inside the vestibule, in particular the FLN6 residues Ala751-Gly778, is seen to interact predominantly with the uL24 loop and H6/H7 region (Fig. 4d and Supplementary Fig. 20). In contrast, the length-equivalent FLN6 residues in FLN5+34 RNCs (Ala751-Val765) interact with H24/H50, as discussed earlier (Fig. 3c, right & Supplementary Fig. 12).

A hypothesis emerges from these cryo-EM data, namely that the structured NC (folded or partially-folded) pathway could be influenced by the stage of biosynthesis (early via H24/H50 and later via H6/H7), where the unstructured NC pathway predominantly explores the H6/H7 region. This was explored further by determination of cryo-EM structures of FLN5+47U RNCs in which FLN5 was rendered folding-incompetent by Y719E mutation^3^. The NC density observed entirely follows the H6/H7 pathway (Fig. 4e) and is continuous from the PTC up to the rRNA H7. *Ab initio* modelling was undertaken and allowed the resolving of a polyalanine model of all but the first four N-terminal FLN6 residues (Ala765-Gly778) before density becomes sparse and not visible at the end of the vestibule just above H7.

### The H6/H7 pathway as a preferred path for unfolded and expanded structures

To complete the picture, we next considered the role of the tunnel at an earlier point in FLN5’s biosynthesis by studying FLN5+31 RNCs, a nascent chain length that is at the cusp of the co-translational folding transition of FLN5^3,18,19^. In solution, FLN5+31 RNCs populate several states^19^, including 44% of the unfolded but also native (∼10%) and intermediate (I1 ∼25% and I2 ∼21%), the latter known to be compact from biophysical analyses and mutational analysis by ^19^F NMR^12,19^. Alongside this system, we studied its variant, FLN5+31 A3A3-RNC (FLN5+31^U^), where FLN5 is entirely disordered^9^ (and therefore expected to follow just one pathway inside the tunnel).

Consistent with the other entirely unfolded NC system from the late stage of biosynthesis (FLN5+47^U^) described above, the entirety of the NC density of FLN5+31^U^ variant was seen following the H6/H7 (Fig.5a). Interestingly, for wild-type FLN5+31-RNC, all 13 classes obtained in its cryo-EM analysis (Fig.5b & Supplementary Fig. 13a,b) again reveal the bifurcation in the NC density close to the uL23 loop (Fig. 5b) with the majority of the observed density following the H24/H50 pathway (Fig.5c, left) as was observed in FLN5+34. NC density consistent with a compact FLN5 domain was again observed only along the H24/H50 pathway (Fig. 5c left), which was constrained within the confines of the vestibule. Comparison between classes that follow one or the other pathway showed that when the NC follows the H24/H50 pathway, hydrogen bonds or/and hydrophobic contacts are formed between A1321 of H50 and C490 of H24 (Fig. 5d), before forming globular structure at the vestibule and beyond. When the NC exits via H6/H7, where density beyond the tunnel is sparse or absent, the NC forms hydrogen bonds and hydrophobic interactions with H6 or H7 and the uL23 loop (Fig. 5d the carbonyl oxygen of FLN5+31 class-9 RNC hydrogen bonds with A73 as well as His70 of rRNA H6 and uL23 loop). The interactions between the ribosome and FLN6 residues involve, (as would be expected) different FLN6 residues to those in FLN5+47 RNCs, which also follow the H6/H7 pathway, and this NC length-dependent network of FLN6-ribosome interactions, as anticipated, suggests a role for these tunnel landmarks in sensing the NC and perhaps steering the NC directionality (Fig. 4b, 5c). Although not the major pathway in FLN5+31, the H6/H7 pathway of FLN5+31 is significantly more populated relative to FLN5+34 and this is in line with the anticipated higher U population present at this shorter length (Fig.4f, Supplementary Fig. 12b). While the density along the H6/H7 pathway is present in several classes, it becomes sparse beyond the vestibule compared to the density within classes following the H24/H50 path (which is more globular). The H6/H7 pathway likely represents the unfolded FLN5 population in this RNC (Fig.5c bottom). These data support the conclusion that the NC can follow two pathways in the tunnel depending on NC length and folding status.

**Figure 5.**
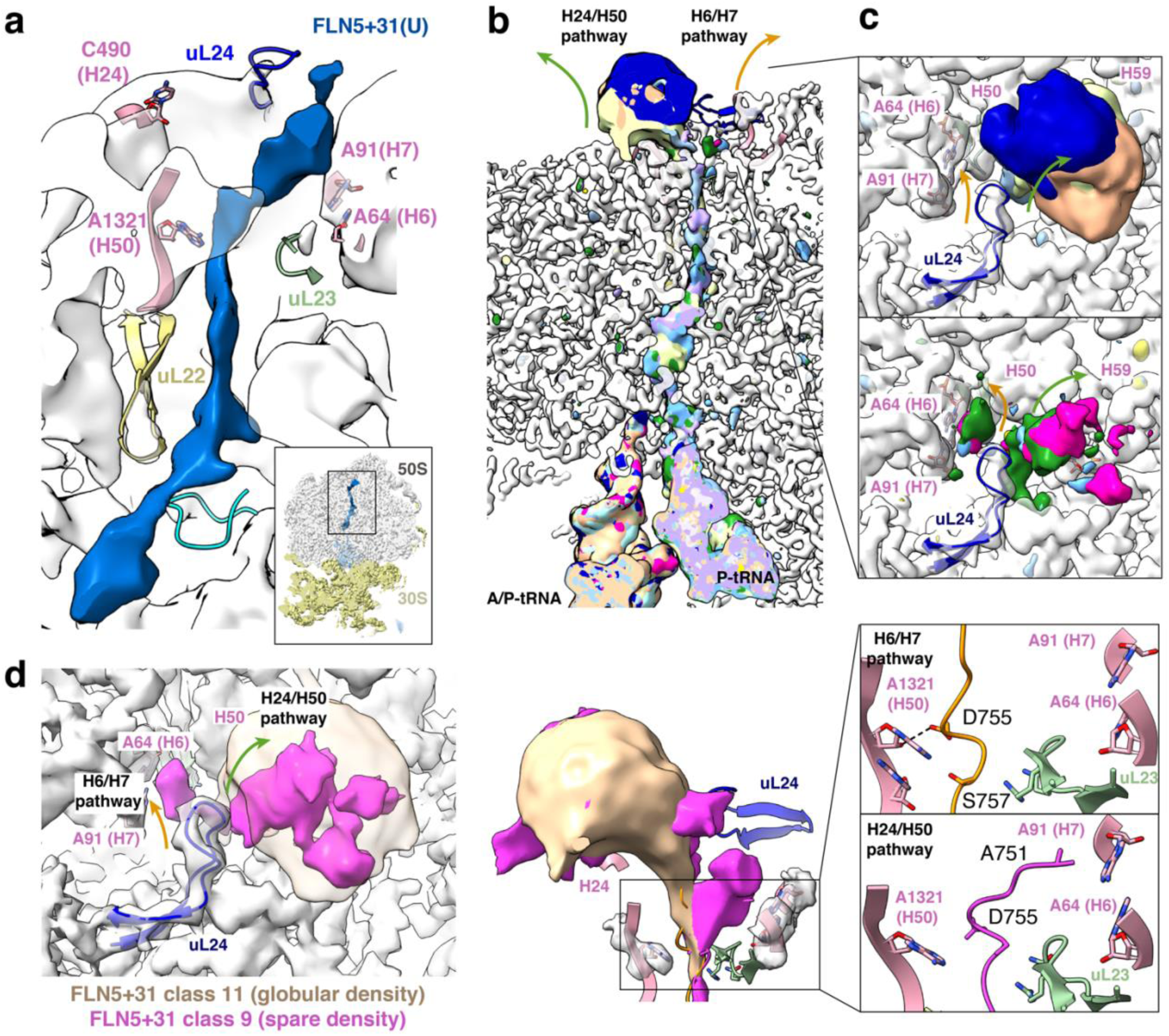
FLN5+31 RNCs show that the H6/H7 pathway is the preferred path for unfolded RNCs. **a.** Reconstruction of the FLN5+31^U^ variant which in solution populates 100% unfolded state. Small and large subunits are shown in yellow and grey surface representation, respectively while the NC is shown in blue. Middle panel: Zoom into the tunnel shows the NC (blue surface) following the H6/H7 pathway at the vestibule. The 50S density is gaussian filtered by 2 and the NC density with sd=1. Right panel: The NC is found near H7 possibly forming interactions/contacts. 50S is shown in white representation and important rRNA bases and ribosomal proteins are noted. **b**. FLN5+31 RNC reconstructions, where folded species are >50% of the population. 13 RNC classes show much of the density following H24/H50 pathway and globular domain is present beyond the vestibule. **c**. Left: FLN5+31 RNC classes following only the H24/H50 pathway. Right: FLN5+31 RNC classes (class-1, −2, −3, 6 and −9) show bifurcation where both pathways are observed. **d**. Comparison between density of class-9 and −11 view from top (left) and transection (right) showing the globular density (class-11) versus sparse density (class-9) and their pathways respectively. Right: close-view of two representative NC models showing H24/H50 pathway and interactions, and H6/H7 pathway and interactions (H6/H7).

To further attempt to understand the observed preferences for unstructured NCs to follow the H6/H7 pathway, we determined the cryo-EM structure of a longer RNC, alpha-1-antitrypsin-RNC (AAT) that our biochemical investigations^45^ suggested would be a useful example: PEGylation studies of the 394 amino acid naturally-stalled AAT-RNC found that the N-terminal region (1-191) of the emerged NC forms a compact protease-resistant molten globule-like state on the ribosome while the C-terminal region of ca. 200 amino acids remains significantly disordered. (Supplementary Fig. 15 for schematic of RNC sequence). The cryo-EM map of SecM-stalled AAT RNCs (Supplementary Fig. 15) shows continuous density inside the tunnel up to the vestibule under the uL24 loop entirely following the H6/H7 pathway. This is in accord with FLN5+47U and FLN5+31 A3A3 RNCs described above where the H6/H7 pathway appears to be a preferred route for NCs that do not have persistent structure within the proximity of the vestibule.

### Elucidation of FLN5 domain conformational heterogeneity and ribosome binding sites

We next considered the interactions between the natively folded state of FLN5 and the ribosome analysed using MD-derived all-atom structural ensembles of FLN5+31, FLN5+34, and FLN5+47 reweighted with the use of cryoENsemble and the cryo-EM maps (see Methods). In the vestibule, for FLN5+31 RNC and FLN5+34 RNCs (Supplementary Fig. 12), the majority of interactions are localised to the uL24 loop and rRNA H24, H50 and H59 with no interactions with the H6/H7 pathway region, while FLN5+47 RNCs interacts almost entirely with the H6/H7 pathway region. We then analysed FLN5’s binding interface to the ribosome surface. In FLN5+31 RNCs, FLN5 occupies a space deep inside the tunnel and thus its interaction interface is found to encompass most of the C-terminal hemisphere of the domain and FLN6 linker (Supplementary Fig. 16b). In FLN5+34 RNCs, ribosome interactions also include the FLN5 C-terminal hemisphere including the A, A’ and G-strands, their corresponding loops as well as part of the FLN6 linker (Supplementary Fig. 16a). Extending the linker by 13 aa in FLN5+47 shows reduced ribosome interactions and a shift in the FLN5-ribosome binding interface towards the N-terminal hemisphere while remaining centred around the G-strand (Supplementary Fig. 16c). The NC binding pattern on the ribosome surface obtained for the reweighted MD structural ensemble recapitulates the pathways observed in the cryo-EM data analysis and provides an atomic-resolution description of these transient binding sites.

### The H6/H7 and H24/H50 NC pathways are present in other RNCs

We compared our observations based on FLN5 RNCs to the other available cryo-EM structures of *E.coli* RNCs. We found regardless of the stalling mode used: SecM^21,27,46^ TnaC^47,48^ VemP^26^, or via a truncated mRNA^49^ that disordered NCs not interacting with any ribosome-associated factors exit the ribosome exclusively via the H6/H7 pathway (Supplementary Fig. 17). These results are thus similar to our pathway preference observations for FLN5 U states described above.

For the globular NC states (of spectrin^32^ and titin^38^) observed within the vestibule the observable density is sparse, suggesting NC conformational heterogeneity in the NC. For spectrin-RNCs the linker density in the vestibule is primarily located close to the H24 and H50 rRNA helices and the globular domain density, which is partially located in the vestibule, is observed to be close to the H24 and uL24 loop. The map also revealed sparse NC density close to H6 and H7 RNA helices, which, in line with the FLN5 observations, suggests that this NC also samples multiple conformational/folding states. The titin I27 domain, on the other hand, exits via a less-defined pathway in the vestibule (Supplementary Fig. 18). This is in generally in line with our recent ^19^F-NMR observations of I27-RNCs of a similar length to cryoEM data presented here (L=34)^12^ which populate several states (native, two intermediates and unfolded states).

Additional published RNC structures containing auxiliary factors, including trigger factor (TF), signal recognition particle (SRP), SecYEG and SecA were also assessed for NC pathway information. Two NC pathways can be traced within the ribosome vestibule for the GatD85 NC obtained with associated TF^50^ from which only the pathway close to the H50 helix directs the NC into the TF cradle (Supplementary Fig. 18). It may be that the unfolded NC, which preferentially interacts with H6 and H7 helices, must shift towards the H50 to be accommodated into the TF cradle. Cryo-EM maps describing different stages of co-translational protein targeting with SRP bound to the ribosome in complex with SR^51,52^, SecA^53^, and SecYEG^51^ all show the NC density following the H24/H50 pathway which presumably is due to the H6/H7 pathway being blocked by the C-terminus of the M domain of the SRP protein (Ffh protein in E. coli) (Supplementary Fig. 19). The H6/H7 pathway is similarly blocked within the complex of the SecYEG translocon-bound ribosome with the 6/7 loop of SecYEG partially inserted into the ribosome exit tunnel (Supplementary Fig. 19). Interestingly, in the case of SecA (another bacterial factor that has a similar role to SRP), the N-terminal tail appears to be too short in length to block the H6/H7 pathway, and the unfolded NC is able to use this path to exit the ribosome (Supplementary Fig. 19).

## Discussion

We set out to investigate the conformational heterogeneity of nascent chains emerging during protein biogenesis, both inside and outside of the ribosomal tunnel, using the immunoglobulin-like domain, FLN5, as our model system. To achieve this, we combined structural analysis by single-particle cryo-EM with Bayesian reweighting of all-atom molecular dynamics simulations. Within this framework, we developed an *in-silico* purification approach for cryo-EM data and applied it to the three stages of FLN5 biosynthesis. This enabled us to visualise the most variable set of NC states on the ribosome to date, comprising of up to 13 classes for each RNC. These classes revealed significant conformational dynamics, highlighting differences not only in the location and orientation of the FLN5 domain on the ribosomal surface but also in the way the NC exits the ribosome through two distinct parallel pathways in the vestibule.

The data described show that the folding status of the nascent polypeptide and its proximity to the exit vestibule as well as any co-factor interactions appear to determine the preferences for the NC within the exit vestibule (Fig. 6). While the H6/H7 pathway is preferred by unfolded NCs, structured NCs prefer the H24/H50 pathway. The latter is likely due to its increased space relative to H6/H7 enabling the NC to fold. The H6/H7 pathway appears to be additionally regulated by the free base pair, A63-A91, from rRNA helices H6 and H7, respectively, as well as the nearby uL23 loop. The H24/H50 pathway, located on the opposite side of the vestibule consists of H50, with A1321 protruding into the tunnel and C490 from H24 pointing to the solvent-exposed region close to the uL24 loop. Both hydrogen bonds and hydrophobic contacts are observed between the nascent polypeptides and these ribosomal elements in both pathways. This points to a potential role of the lower tunnel components, and more specifically rRNA helices H6, H7, H24 and H50, in pathway selection and, hence, co-translational folding and perhaps even targeting. By removing crucial interactions with the RNA helices H6 and H7 (using FLN5+34_23i)^30^, we have demonstrated how one pathway can be blocked and affect co-translational folding. Ribosome-associated factors appear to be involved in the further modifying the co-translational pathway (TF, SRP, SecY). With trigger factor (TF) present, cryo-EM shows the unfolded GatD85 RNC^50^ to follow two pathways, one along the ‘canonical’ H6/H7 route and the other directing the NC towards the TF cradle.

**Figure 6.**
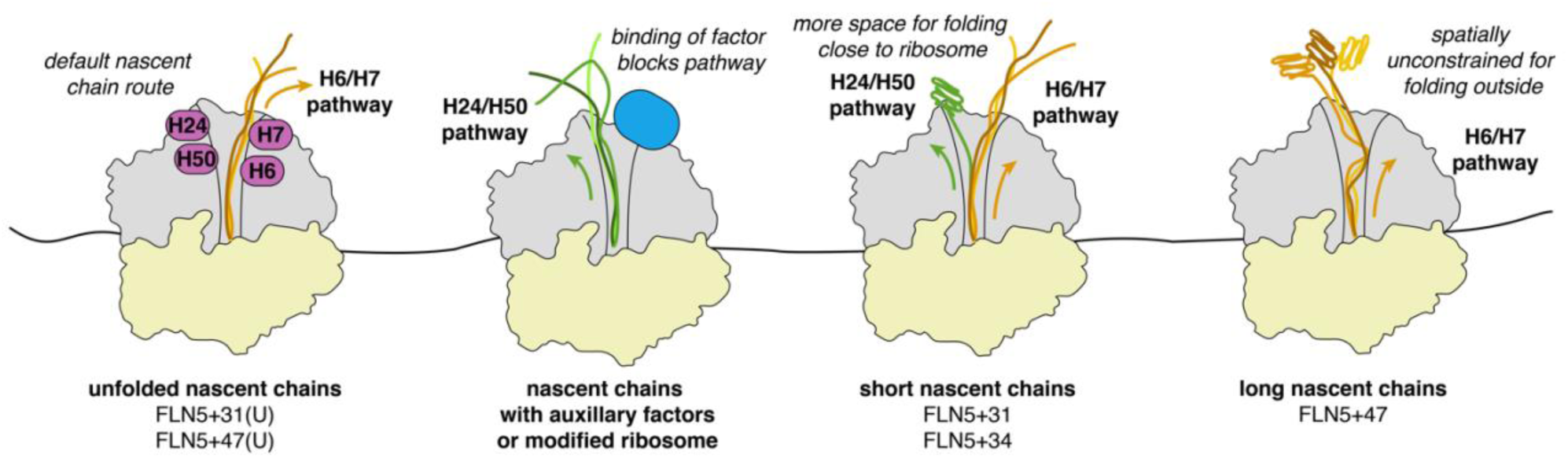
Summary of structural biosynthetic snapshots of FLN5 during the folding transition. For unfolded nascent chains (folding-incompetent mutants at a short (L=31) and longer (L=47) linker length) where only the U state is visible with NMR, we visualise cryo-EM density that follows pathway H24/H50. At short length (L=31 and L=34) two pathways are visible (purple) and the cryo-EM density corresponding to the folded FLN5 domain can be seen partially in (L=31) or near (L=34) the vestibule (purple box) when the NC follows pathway H24/H50. At longer linker length (FLN5+47), all states follow pathway H6/H7 in the vestibule, there are several globular densities (for FLN5+47) similar to the shorter lengths but at longer distance from the ribosomal surface Ribosome is shown in grey surface, the tunnel wall is shown in black lines. The position of H6-H7 free base pair and H50/H24 is highlighted in all schematic representation.

The integrative structural biology approach has allowed us to characterise FLN5 RNC structural heterogeneity, including the description of the interactions between the FLN5 domain and the ribosome. The emerging picture from these analyses shows that the folded nascent chain is highly dynamic, sampling multiple transient binding sites. This is supported by our simulations of FLN5+47 that are in good agreement with both cryo-EM maps (Supplementary Fig. 7) and NMR measurements of rotational tumbling times (Supplementary Fig. 5). This work thus forms the basis for future integrative structural studies on the biosynthesis of specific proteins and their exit and folding paths from the vestibule, including their interactions with chaperones and other factors involved in NC processing.

## Methods

### Generation of ribosomal nascent chain complexes (RNCs) for cryo electron microscopy

All SecM-stalled RNC constructs of FLN5 (L=31, 31U, 34, 47 and 47U) were generated as detailed previously^3,18^. The enhanced translational arrest SecM sequence was used for all RNCs and the RNCs were generated in BL21(DE3) E. coli as described previously^3^. The FLN5 RNC at L=34 was generated in CRISPR-modified BL21(DE3) E. coli cells where the uL23 was modified to have a longer loop (insertion) in the tunnel as previously described^30^. The cells were lysed with French press and purification was performed via sucrose cushion sedimentation, Nickel affinity and butyl chromatography as previously described. The pooled fractions were concentrated with centrifugal filter units at 3,000g, with buffer changed to EM buffer. All RNCs were resuspended in EM buffer and the sample concentrations measured by absorbance in 260nm and the 260/280 was monitored to be >1.9. The purity and occupancy of the RNC samples was monitored with low-pH gel electrophoresis and immunoblotting for visualisation by anti-HisTag western blotting. The RNC was separated via semi-denaturing page and transferred to nitrocellulose blotting membrane. After blocking with 5% skim milk, the western blots were incubated with anti-His antibody overnight and developed with pico chemiluminescent substrate.

### Cryo-EM grid preparation and data acquisition

#### FLN5+34

3μl of 250nM purified RNC were applied onto glow-discharged (30s) holey carbon grids pre-coated with 2nm carbon (Quantifoil 2/2) layer using Vitrobot with the following parameters; humidity 95%, temperature 4°C, incubation time 30s, blotting force −10 and blotting time for 9s from both sides. Data acquisition was performed at eBIC on Titan Krios IV equipped with K3 camera using EPU and voltage 300keV. Four shots per holes were acquired with faster acquisition 295 images/hour and an energy slit with width of 20keV was used during data collection. 11,502 movies at super resolution pixel size 0.536Å were collected at defocus range 1.2-2.4μm. The dose per frame was ∼1.08 e^-^/Å2 and the 40 frames were aligned using Motioncorr with dose-weighting in Relion 4.0 on-the-fly.

#### FLN5+47

Freshly purified FLN5+47 sample was diluted to 50nM in EM buffer and 3μL were applied at 4°C and 100% humidity onto glow discharged Quantifoil R1.2/1.3 grids coated with in-house prepared graphene oxide. After incubation for 10s and blotting for 11s the grid was vitrified into liquid ethane using Vitrobot. Both datasets were collected at Titan Krios TEM equipped with K2 detector at 300keV under low dose conditions. 10,177 movies were collected at super-resolution pixel size was 0.5335 and with a defocus range of 0.5-2.5 um. The dose was 0.97 e^-^/Å2 per frame and the 50 frames were aligned using Motioncorr in Relion 3.1^54^.

#### FLN5+31

Glow-discharged holey carbon grids pre-coated with 2nm carbon (Quantifoil 2/2) layer were used to apply 3μl of purified RNC using Vitrobot (humidity 95%, temperature 4°C, incubation time 30s, blotting force −10 and blotting time for 8.5s from both sides). Grids were transferred into a Titan Krios II (eBIC) electron microscope (Thermo Fisher Scientific) operating at 300 keV and data acquired with a K3 camera with super resolution mode bin 2 (Gatan, Inc.). The ∼38,000 image stacks (movies) were acquired with a pixel size of 0.831 Å/pixel using the EPU software (ThermoFisher Scientific) to record movies with 50 fractions with a total accumulated dose of ∼46 e^-^/Å2/movie. The defocus values ranging between –0.2 and –2.1 μm.

#### FLN5+47U

3μl of 160nM of purified RNC was applied to freshly glow-discharged Quantifoil 3/3 holey carbon grids pre-coated with 3nm carbon. Vitrification was performed with 8s blotting, blot force −10 and the vitrobot chamber at 100% humidity and 4C. Data acquisition was performed in Titan Krios equipped with K2 camera in counting mode. 5,500 images were collected with a pixel size of 1.05 Å/px using EPU software to record movies with 32 fractions with a total accumulated dose of ∼ e^-^/Å2/movie and defocus range −0.6 to 3μm.

#### FLN5+31U

5μl of 125nM purified FLN5+31U RNC were applied to glow-discharged holey carbon grids pre-coated with 2nm carbon (Quantifoil 2/2) layer and vitrified using Vitrobot (humidity 95%, temperature 4°C, incubation time 30s, blotting force −10 and blotting time for 8.5s from both sides). Grids were transferred into Titan Krios III (eBIC) electron microscope (Thermo Fisher Scientific) operating at 300 keV This data was acquired with TFS Selectris X and Falcon 4i 4096 x 4096. The ∼38,000 image stacks (movies) were acquired with a pixel size of 0.723/2 Å/pixel using the EPU software (ThermoFisher Scientific) to record movies with 50 frames with 1e^-^ per frame, with a total accumulated dose of ∼50 e^−^/Å^2^/movie and total EER fraction of ∼18 to sum per frame. The defocus values ranging between –0.2 and –2.1 μm.

### Cryo-EM image processing

For FLN5+34 pre-processing (motion correction and CTF estimation) of the 11,502 micrographs was performed in Relion 4.0^42^. After visual inspection for optimal ice quality, absence of astigmatism and/or drift, as well as selection for resolution between that 3Å the selected 10,803 micrographs after were subjected to topaz picking and yielded to 5,322,230. The particles, downsized by a factor of four, was subjected to a reference-free 2D classification. After visual inspection of the 2D classes the 2,592,748 particles were 3D refined to a reconstruction using a 70S ribosome as reference with low-pass filtering at 50Å. Further heterogeneity of these particles was treated as shown in Supplementary Fig.2: briefly, the initial reconstruction was subjected to 3D classification with a mask in the intersubunit space and the states isolated were A/A and P/P and very weak E/E tRNAs (AP-tRNA), A/A and P/P and E/E tRNA (APEtRNA) and a state with a P-tRNA only. The two additional classes showed 70S with no apparent density for tRNAs or density corresponding to the 50S subunit. The AP-tRNA class was subjected in further 3D classification with mask around the EtRNA to remove particles with E/E tRNA only and thus no NC present. The AP-tRNA and P-tRNA RNCs were refined with a 50S mask and the re-extracted particles (not downsized) were reextracted and the images refined before CTF refinement and Bayesian particle polishing. At this point an additional round of 3D classification was used with an ellipsoid soft-edged mask covering the tunnel as well as density for the rRNA and ribosomal proteins surrounding the tunnel (see mask details below). The particles were sorted in six classes for the P-tRNA and AP-tRNA RNCs, respectively. Fourier shell correlation (FSC) calculation between independently refined random half subsets, local resolution estimation and post-processing were done in RELION 4.0^42^.

Similar data processing pipeline was used for FLN5+47 RNC. Briefly, CTFFIND-4.1 was used to determine power spectra, defocus values and astigmatism. After manual inspection the micrographs were filtered by threshold of figure-of-merit 0.15 and resolution at 4.5Å resulting in 9,295 micrographs. 995,570 particles were picked by Gautomatch (external job in Relion 3.1^54^) and subjected to 2D classification. 648,915 particles were subjected to 3D refinement using E.coli ribosome as reference map and low-pass filtered it at 60Å. After 3D classification 260,662 particles with Psite present in unratcheted ribosomes were selected and after 3D refinement the particles were subjected in focused 3D classification with an ellipsoid mask at the intersubunit space covering A-, P- and E-tRNA sites. Three major classes containing 77,590, 100,252 and 84,820 particles were further refined into P-tRNA RNC, AP-tRNAs RNC and empty ribosomes respectively. The P-tRNA RNC and AP-tRNAs RNC reconstructions were further subjected in postprocessing and CTF refinement for aberrations, magnification and defocus. The resulted particles were polished using Bayesian Polishing and the resolution for the P-tRNA RNC and AP-tRNAs RNC reconstructions was 2.55 and 2.45 Å, respectively. The P-tRNA RNC particles were subjected in an extra round of 3D classification with a mask including most of the tunnel and 40 Å of the space outside the vestibule. This resulted into four and six major classes for the P-tRNA RNC and AP-tRNAs RNC, respectively. Each class was refined and the local resolution was estimated using ResMap. The AP-tRNA RNC particles were subjected in a similar round of 3D classification with a mask including most of the tunnel and 40 Å of the space outside the vestibule. This classification resulted in 3 classes two of which had clear density on the outside (classes 1 and 2) whereas the highest populated class (class-3) had no density for the globular domain. The latter was subjected to an extra round of focused classification revealing four distinct conformations for the globular domain (classes 3-6 supp figure 1).

For FLN5+47U pre-processing (motion correction and CTF estimation) of the 5,500 micrographs was performed in Relion 4.0^42^. After visual inspection the selected micrographs were subjected to topaz picking and yielded to 589,792 particles. The particles, downsized by a factor of four, was subjected to a reference-free 2D classification. After visual inspection of the 2D classes the 497,151 particles were 3D refined to a reconstruction using a 70S ribosome as reference with low-pass filtering at 50Å. After two rounds of 3D classifications with and without mask in the intersubunit space 143,483 particles were extracted in normal size (unbinned). 3D refinement of these particles yielded a map of 2.6Å. The refined particles were subjected to per particles defocus, magnification and astigmatism CTF refinement and Bayesian polishing.

For FLN5+31, following quality control inspection, 1,206,843 particles were picked with Topaz particle picking from 10,167 movie stacks and subjected to reference-free 2D classification. The four times downsized particles were refined and used for structural heterogeneity as follows (Supplementary Fig. 14). As with FLN5+34 the initial dataset was subjected to 3D classification with mask in the intersubunit space and refined without binning. The refined particles were subjected to per particles defocus, magnification and astigmatism CTF refinement and Bayesian polishing before being subjected to tunnel 3D classifications.

For FLN5+31U, which was collected with Falcon 4i, we used similar approach to the datasets above. After quality inspection 27,465 micrographs were selected. Topaz picking yielded to an average of 144 particles per micrographs and after setting the autopick FOM threshold at −3, 3,616,234 particles four-times downscaled were extracted. Inspection of the 2D classes yielded to 2,635,500 particles which were 3D refined and 3D classified without mask. The particles with P-site present (1894401) were extracted in original pixel size and the resolution of the 3D refined map was 2.7 Å. Local resolution variability (Supplementary Fig. 3,11) was estimated via ResMap or Relion local resolution and visualized by coloring the corresponding maps in ChimeraX^55^.

### Density model for ellipsoid mask used for 3D classifications

For generation of the constant density volume for the tunnel mask for 3D classifications we used SPIDER. Using the MO operation we created a custom-made mask to include the tunnel from the PTC to the uL24 and beyond with x,y,z radii (std.dev.=2,3,4) (Fig. 2a). This mask was then resampled on the grid dimensions of the 3D refined map and the output map was then input in Relion Mask Creation tool with initial binarization 0.1, addition of soft edge by 6pixels and extended the binary map by 3pixels. The choice of extending the mask beyond the vestibule but not extensively is related to the dynamic nature of the globular domain and the limitations of cryo-EM. For such a dynamic system to be visible with cryo-EM, a highly populated bound state should be captured, which should be near the ribosomal surface.

### Model building

The 6qdw E.coli 70S ribosome without tRNAs and NC was used for the molecular modeling of the 70S ribosome. The proline tRNA was used from the Pro-tRNA of the TnaC model and rigid-body-fitted into the cryo-EM densities using ChimeraX^55^, followed by iterative refinement using PHENIX^56^ real-space refinement and manual adjustment in Coot^57^. The AE1 was built denovo for the highest resolution reconstruction (class_4) and residues from the linker were also adjusted manually. For the rest of the conformations the class-4 AE1 model was rigid body fitted and adjusted where needed. To help with model building and only, maps were modified using Resolve in PHENIX, where necessary. The de novo model building was done in 3 σ for the AE1 and at 1.5-2.8 σ for the FLN6 linker.

For the model building of the FLN5 globular domain a map was generated from the NMR structureσ from the full length FLN5 (pdb:6G4A) or the x-ray structure^58^ of the full-length FLN5 and it was subsequently fitted into the isolated FLN5 globular density of each classes. The fit with the highest Fourier-shell correlation was the best orientation possible for each conformation.

### Model analysis

Molrpobity was used to calculate the statistics of the refined models. FSC curves between the model and the map were also calculated for validation RMSD for the 50S was performed between the LSU of P-site or AP-site before tunnel classification. RMSDs between different NCs were calculated in ChimeraX^55^ and Pymol and are indicated in brackets in the main text next to relevant sections.

### Setting up all-atom molecular dynamics simulations

We performed all-atom MD simulations of the FLN5 folded state on the ribosome at linker lengths of FLN5+31, FLN5+34 and FLN5+47 with GROMACS 2021^59^ using the CHARMM36m force field and CHARMM-modified TIP3P (cTIP3P) water model^60^. Initial simulations of FLN5+47 were performed with the original cTIP3P parameters (termed C36m here), leading to overstabilisation of FLN5 binding to the ribosome surface compared to NMR data (Supplementary Fig. 5g). We thus, following^60^, decreased the hydrogen Lennard-Jones ε parameter of cTIP3P water (termed C36m+W here) to −0.1 kcal mol^-1^ (instead of the original −0.046 kcal mol^-1^), which resulted in reduced FLN5-ribosome interactions and improved agreement with NMR measurements of the rotational correlation time of FLN5 (Supplementary Fig. 5g).

To build starting models for FLN5 RNCs MD simulations, we used, as in our previous work^9^, a model of the ribosomal exit tunnel with surrounding surface generated based on the high-resolution 70S E. coli ribosome structure (PDB id: 4YBB^61^). The initial structure of the FLN6-SecM linker was obtained from the FLN5+47 RNC model before vestibule classification, which was then used to also model the other lengths (+31, +34 RNC). We attached the folded structure of FLN5 (PDB 1QFH^58^) to the N-terminus of the linker and added the disordered structure of the His-tag to the N-terminus of FLN5. We used a dodecahedron simulation box and solvated and neutralised the system with cTIP3P water and Mg^2+^ ions. This resulted in final system sizes of 1,461,410, 1,461,577 and 1,461,380 atoms with 705 Mg^2+^ (FLN5+47) or 706 Mg^2+^ (FLN5+31 and FLN5+34) ions. Energy minimisation of the systems was performed with the steepest-decent algorithm. For equilibration and production runs the LINCS algorithm^62^ was used to constrain all bonds involving hydrogen. Equilibration runs were performed with a 2 fs timestep and van der Waals interactions were treated with a 1.2 nm cut-off and a switching function from 1.0 nm. Real-space electrostatic interactions were calculated up to a cut-off of 1.2 nm, and long-range interactions were treated with the Particle Mesh Ewald (PME) method^63^. The systems were first equilibrated in the NVT ensemble for 1 ns at a temperature of 298 K using the velocity rescaling algorithm^64^ with a time constant of 0.1 ps and with position restraints assigned to all heavy atoms (i.e., the NC and ribosome atoms) with a force constant of 1000 kJ mol^-1^ nm^-2^. Then, the system was equilibrated in the NPT ensemble for 1 ns at 1 bar with a compressibility of 4.5×10-5 bar^-1^ using the Berendsen barostat^65^, followed by another 1 ns simulation with the Parrinello-Rahman algorithm^66^. For subsequent production simulations in the NPT ensemble, heavy atom position restraints were kept for all ribosome heavy atoms and the C-terminal NC residue at the PTC but removed for the rest of the NC. We then ran ∼400 ns of unbiased MD to generate different starting structures for final production simulations. Four production simulations from different starting structures were run for 2 µs each with hydrogen mass repartitioning applied^67^ (hydrogen masses scaled by a factor of 4), allowing for an integration timestep of 4 fs. Coordinates of the RNCs were saved every 80 ps for analysis, resulting in a total of 100,000 frames from the four, concatenated trajectories per RNC. FLN5 remained folded in all simulations.

The obtained MD trajectories represented a diverse set of NC conformations (Supplementary Fig. 17), which we further clustered using the gromos clustering method implemented in GROMACS, selecting a cut-off (1.0 nm for +47 RNC) that can give us ∼1000 clusters. Centroid structures from each cluster were combined to form prior MD ensembles of FLN5+31, FLN5+34, and FLN5+47 RNCs (Schrödinger LLC, version 2.4.2) reweighting. This number of structures is computationally manageable and provides an excellent structural representation of the trajectory for subsequent reweighting in cryoENsemble.

### Comparison of molecular dynamics simulation with NMR data

To compare our simulations with NMR data reporting on the strength of nascent chain-ribosome interactions, we calculated the rotational correlation time of FLN5 as this property is highly sensitive to ribosome interactions^34^. Because simulations employing the TIP3P water model do not result in accurate predictions of absolute solute and solvent diffusion and rotational properties^68^, we used MD simulations of the isolated protein to derive a normalised value to compare with experiments. Isolated FLN5 simulations (residues M637 to G750, initiated from the crystal structure PDB code:1QFH) were performed using an identical protocol as detailed above for the RNC. A large box size was chosen with FLN5 placed at least 2 nm from the closest edge. The box volume was 916 nm^3^ and contained 29,384 water molecules. To calculate the rotational correlation time of FLN5, we utilised 1,000 vectors uniformly distributed in the unit sphere. Heavy backbone atoms of the native β-strands were used to fit the domain and rotation matrices were applied to all vectors in the unit sphere and correlation function was then calculated using a second-order Legendre polynomial^69^:

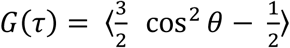

θ is the angle between a vector at time t and t+τ after rotation. We averaged over all pairs of frames separated by the lag time τ. Data of all vectors up to lag times of 50 ns were then fit globally to a single exponential to determine the rotational correlation time, τC.

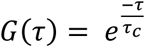

We used four simulations of 2 μs to determine the mean τC and standard error of the mean. For RNCs, data were fit to a double exponential and a weighted fit to the average and standard deviation obtained from all 1,000 vectors.

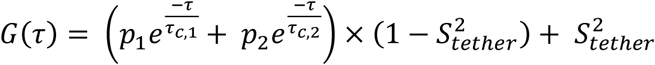

Here, p1 and p2 correspond to the populations/amplitude of the two exponential decays and S^2^tether is an order parameter describing the extent to which the position-restrained ribosome restricts FLN5 tumbling. For correlation functions that decay to 0, S^2^tether is set to 0. The four independent MD trajectories were fit individually to obtain the mean τC and standard error of the mean. For each trajectory, the time-averaged τC was obtained as τ_*c*_ = (*p*_1_τ_*c*,1_ + *p*_2_τ_*c*,2_) × (1 − *S*_*tether*_^2^) + *S*_*tether*_^2^ τ_*r*,70*S*_ where τ_*r*,70*S*_ is the ribosome rotational correlation time under these conditions (∼3,000 ns)^70^. Since the TIP3P water model results in faster protein/FLN5 correlation times, however, we scaled τ_*r*,70*S*_ according to the factor by which protein tumbling is attenuated relative to experiments (in isolation, 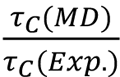
).

For reweighting with cryo-EM maps, the MD trajectories represented a diverse set of NC conformations Supplementary Fig. 17), which we further clustered using the gromos clustering method implemented in GROMACS, selecting a cut-off (1.0 nm for +47 RNC) that can give us ∼1000 clusters. Centroid structures from each cluster were combined to form prior MD ensembles of +31, +34, and +47 RNCs (Schrödinger LLC, version 2.4.2) reweighting. This number of structures is computationally manageable and provides an excellent structural representation of the trajectory.

### The cryoENsemble reweighting of the MD-derived structural ensemble

For the reweighting, we selected classes with cryo-EM maps that show a clear density outside the vestibule, at the 3σ threshold (for FLN5+47 and FLN5+34 RNCs) and 2σ threshold (for FLN5+31 RNC), that can be attributed to the structured FLN5. This selection resulted in 23 datasets, including nine classes for FLN5+31 (classes-1,3,4,5,6,8,10,11 and 12), ten classes for FLN5+34 (classes-1, 2, 5-12) and four classes for FLN5+47 (classes-2, 5, 6, 9). To obtain cryo-EM density that corresponds only to the NC, we subtracted the empty ribosome density from each RNC’s cryo-EM map using the volume zone and volume subtract commands in ChimeraX^55^ to obtain cryo-EM density that corresponds only to the NC. Using ChimeraX, we applied the volume gaussian command with 3Å standard deviation to each NC map to smooth the map and improve the signal-to-noise ratio, and we binned the map with volume bin command to decrease the voxel size to ∼2Å, which significantly reduces the number of voxels for reweighting while preserving the information.

Prior to the reweighting, we fitted NC structural ensembles into the cryo-EM maps by structurally aligning each RNC structure to the ribosome structure pre-fitted to the cryo-EM map in ChimeraX^55^, using only the ribosome structure as a reference. We applied cryoENsemble following the iterative methodology described in Wlodarski et al^71^ (Supplementary Fig. 5) to generate new posterior NC structural sub-ensembles (Supplementary Fig.18-20) for each selected NC cryo-EM map. All the subensembles had improved agreement with the corresponding NC cryo-EM maps, which we assessed based on the calculated correlation coefficient (Supplementary Fig. 19).

## Data availability

The coordinates are deposited in the Protein Data Bank (PDB) with accession codes PDB xxxx. The cryo-EM maps have been deposited in the Electron Microscopy Data Bank (EMDB) with accession codes EMD-xxxxx. Other relevant data generated in this study are provided in the Supplementary Information.

## Acknowledgements

We acknowledge ISMB EM facility (Birkbeck College, University of London) with financial support from the Wellcome Trust (202679/Z/16/Z and 206166/Z/17/Z). We thank Minkoo Ahn for help with protein production. We thank Natasha Lukoyanova and Shu Chen for data collection and David Houldershaw for help with computing in data processing. We acknowledge Diamond for access and support of the cryo-EM facilities at the UK national electron Bio-Imaging Centre (eBIC), proposals em20287, bi26703 and bi34130 funded by the Wellcome Trust, Medical Research Council and BBSRC. This project made use of time on HPC resources on Archer2 (ARCHER2 UK National Supercomputing service, https://www.archer2.ac.uk) granted via the UK High-End Computing Consortium for Biomolecular Simulation, HECBioSim (http://hecbiosim.ac.uk), supported by EPSRC (grant no. EP/R029407/1 and EP/X035603/1). We also acknowledge the EuroHPC Joint Undertaking for awarding this project access to the EuroHPC supercomputer LUMI, hosted by CSC (Finland) and the LUMI consortium through a EuroHPC Regular Access call and the Baskerville Tier 2 HPC service (https://www.baskerville.ac.uk/). Baskerville was funded by the EPSRC and UKRI through the World Class Labs scheme (EP/T022221/1) and the Digital Research Infrastructure programme (EP/W032244/1) and is operated by Advanced Research Computing at the University of Birmingham. We additionally acknowledge the use of the UCL Myriad and Kathleen High Performance Computing Facility (Myriad@UCL and Kathleen@UCL), and associated support services, in the completion of this work.

## References

1 Cassaignau, A. M. E., Cabrita, L. D. & Christodoulou, J. How Does the Ribosome Fold the Proteome? Annu Rev Biochem 89, 389–415 (2020). 10.1146/annurev-biochem-062917-012226

2 Jumper, J. et al. Highly accurate protein structure prediction with AlphaFold. Nature 596, 583–589 (2021). 10.1038/s41586-021-03819-2

3 Cabrita, L. D. et al. A structural ensemble of a ribosome-nascent chain complex during cotranslational protein folding. Nat Struct Mol Biol 23, 278–285 (2016). 10.1038/nsmb.3182

4 Waudby, C. A., Dobson, C. M. & Christodoulou, J. Nature and Regulation of Protein Folding on the Ribosome. Trends Biochem Sci 44, 914–926 (2019). 10.1016/j.tibs.2019.06.008

5 Holtkamp, W. et al. Cotranslational protein folding on the ribosome monitored in real time. Science 350, 1104–1107 (2015). 10.1126/science.aad0344

6 Kemp, G., Kudva, R., de la Rosa, A. & von Heijne, G. Force-Profile Analysis of the Cotranslational Folding of HemK and Filamin Domains: Comparison of Biochemical and Biophysical Folding Assays. J Mol Biol 431, 1308–1314 (2019). 10.1016/j.jmb.2019.01.043

7 Liutkute, M., Maiti, M., Samatova, E., Enderlein, J. & Rodnina, M. V. Gradual compaction of the nascent peptide during cotranslational folding on the ribosome. Elife 9 (2020). 10.7554/eLife.60895

8 Notari, L., Martinez-Carranza, M., Farias-Rico, J. A., Stenmark, P. & von Heijne, G. Cotranslational Folding of a Pentarepeat beta-Helix Protein. J Mol Biol 430, 5196–5206 (2018). 10.1016/j.jmb.2018.10.016

9 Cassaignau, A. M. E. et al. Interactions between nascent proteins and the ribosome surface inhibit co-translational folding. Nat Chem 13, 1214–1220 (2021). 10.1038/s41557-021-00796-x

10 Kaiser, C. M., Goldman, D. H., Chodera, J. D., Tinoco, I., Jr. & Bustamante, C. The ribosome modulates nascent protein folding. Science 334, 1723–1727 (2011). 10.1126/science.1209740

11 Deckert, A. et al. Common sequence motifs of nascent chains engage the ribosome surface and trigger factor. Proc Natl Acad Sci U S A 118 (2021). 10.1073/pnas.2103015118

12 Streit, J. O. et al. The ribosome lowers the entropic penalty of protein folding. Nature (2024). 10.1038/s41586-024-07784-4

13 Akbar, S., Bhakta, S. & Sengupta, J. Structural insights into the interplay of protein biogenesis factors with the 70S ribosome. Structure 29, 755–767 e754 (2021). 10.1016/j.str.2021.03.005

14 Bhakta, S., Akbar, S. & Sengupta, J. Cryo-EM Structures Reveal Relocalization of MetAP in the Presence of Other Protein Biogenesis Factors at the Ribosomal Tunnel Exit. J Mol Biol 431, 1426–1439 (2019). 10.1016/j.jmb.2019.02.002

15 Cabrita, L. D., Dobson, C. M. & Christodoulou, J. Protein folding on the ribosome. Curr Opin Struct Biol 20, 33–45 (2010). 10.1016/j.sbi.2010.01.005

16 Hartl, F. U. & Hayer-Hartl, M. Converging concepts of protein folding in vitro and in vivo. Nat Struct Mol Biol 16, 574–581 (2009). 10.1038/nsmb.1591

17 Schwarz, A. & Beck, M. The Benefits of Cotranslational Assembly: A Structural Perspective. Trends Cell Biol 29, 791–803 (2019). 10.1016/j.tcb.2019.07.006

18 Waudby, C. A. et al. Systematic mapping of free energy landscapes of a growing filamin domain during biosynthesis. Proc Natl Acad Sci U S A 115, 9744–9749 (2018). 10.1073/pnas.1716252115

19 Chan, S. H. S. et al. The ribosome stabilizes partially folded intermediates of a nascent multi-domain protein. Nat Chem 14, 1165–1173 (2022). 10.1038/s41557-022-01004-0

20 Waudby, C. A., Burridge, C., Cabrita, L. D. & Christodoulou, J. Thermodynamics of co-translational folding and ribosome-nascent chain interactions. Curr Opin Struct Biol 74, 102357 (2022). 10.1016/j.sbi.2022.102357

21 Bhushan, S. et al. SecM-stalled ribosomes adopt an altered geometry at the peptidyl transferase center. PLoS Biol 9, e1000581 (2011). 10.1371/journal.pbio.1000581

22 Bischoff, L., Berninghausen, O. & Beckmann, R. Molecular basis for the ribosome functioning as an L-tryptophan sensor. Cell Rep 9, 469–475 (2014). 10.1016/j.celrep.2014.09.011

23. Gersteuer, F., et al. Publisher Correction: The SecM arrest peptide traps a pre-peptide bond formation state of the ribosome. Nat Commun 15, 3276 (2024). 10.1038/s41467-024-47509-9

24. Morici, M., et al. Publisher Correction: RAPP-containing arrest peptides induce translational stalling by short circuiting the ribosomal peptidyltransferase activity. Nat Commun 15, 3242 (2024). 10.1038/s41467-024-47508-w

25 Sohmen, D. et al. Structure of the Bacillus subtilis 70S ribosome reveals the basis for species-specific stalling. Nat Commun 6, 6941 (2015). 10.1038/ncomms7941

26 Su, T. et al. The force-sensing peptide VemP employs extreme compaction and secondary structure formation to induce ribosomal stalling. Elife 6 (2017). 10.7554/eLife.25642

27 Zhang, J. et al. Mechanisms of ribosome stalling by SecM at multiple elongation steps. Elife 4 (2015). 10.7554/eLife.09684

28 Nilsson, O. B. et al. Cotranslational Protein Folding inside the Ribosome Exit Tunnel. Cell Rep 12, 1533–1540 (2015). 10.1016/j.celrep.2015.07.065

29 Agirrezabala, X. et al. A switch from alpha-helical to beta-strand conformation during co-translational protein folding. EMBO J 41, e109175 (2022). 10.15252/embj.2021109175

30 Ahn, M. et al. Modulating co-translational protein folding by rational design and ribosome engineering. Nat Commun 13, 4243 (2022). 10.1038/s41467-022-31906-z

31 Kemp, G., Nilsson, O. B., Tian, P., Best, R. B. & von Heijne, G. Cotranslational folding cooperativity of contiguous domains of alpha-spectrin. Proc Natl Acad Sci U S A 117, 14119–14126 (2020). 10.1073/pnas.1909683117

32 Nilsson, O. B. et al. Cotranslational folding of spectrin domains via partially structured states. Nat Struct Mol Biol 24, 221–225 (2017). 10.1038/nsmb.3355

33 Streit, J. O. et al. Long-range electrostatic forces govern how proteins fold on the ribosome. bioRxiv, 2025.2002.2010.637539 (2025). 10.1101/2025.02.10.637539

34 Burridge, C. et al. Nascent chain dynamics and ribosome interactions within folded ribosome-nascent chain complexes observed by NMR spectroscopy. Chem Sci 12, 13120–13126 (2021). 10.1039/d1sc04313g

35 Pellowe, G. A. et al. The human ribosome modulates multidomain protein biogenesis by delaying cotranslational domain docking. bioRxiv, 2024.2009.2019.613857 (2024). 10.1101/2024.09.19.613857

36 Roeselova, A. et al. Mechanism of chaperone coordination during cotranslational protein folding in bacteria. Mol Cell 84, 2455–2471 e2458 (2024). 10.1016/j.molcel.2024.06.002

37 Wales, T. E. et al. Resolving chaperone-assisted protein folding on the ribosome at the peptide level. Nat Struct Mol Biol 31, 1888–1897 (2024). 10.1038/s41594-024-01355-x

38 Tian, P. et al. Folding pathway of an Ig domain is conserved on and off the ribosome. Proc Natl Acad Sci U S A 115, E11284–E11293 (2018). 10.1073/pnas.1810523115

39 Cassaignau, A. M. et al. A strategy for co-translational folding studies of ribosome-bound nascent chain complexes using NMR spectroscopy. Nat Protoc 11, 1492–1507 (2016). 10.1038/nprot.2016.101

40 Yu, S., Srebnik, S. & Dao Duc, K. Geometric differences in the ribosome exit tunnel impact the escape of small nascent proteins. Biophys J 122, 20–29 (2023). 10.1016/j.bpj.2022.11.2945

41 Włodarski, T. et al. CryoENsemble - a Bayesian approach for reweighting biomolecular structural ensembles using heterogeneous cryo-EM maps. bioRxiv, 2023.2011.2021.567999 (2023). 10.1101/2023.11.21.567999

42 Zivanov, J. et al. A Bayesian approach to single-particle electron cryo-tomography in RELION-4.0. Elife 11 (2022). 10.7554/eLife.83724

43 Punjani, A., Rubinstein, J. L., Fleet, D. J. & Brubaker, M. A. cryoSPARC: algorithms for rapid unsupervised cryo-EM structure determination. Nat Methods 14, 290–296 (2017). 10.1038/nmeth.4169

44 Zhong, E. D., Bepler, T., Berger, B. & Davis, J. H. CryoDRGN: reconstruction of heterogeneous cryo-EM structures using neural networks. Nat Methods 18, 176–185 (2021). 10.1038/s41592-020-01049-4

45 Plessa, E. et al. Nascent chains can form co-translational folding intermediates that promote post-translational folding outcomes in a disease-causing protein. Nat Commun 12, 6447 (2021). 10.1038/s41467-021-26531-1

46 Schulte, L. et al. Cysteine oxidation and disulfide formation in the ribosomal exit tunnel. Nat Commun 11, 5569 (2020). 10.1038/s41467-020-19372-x

47 Seidelt, B. et al. Structural insight into nascent polypeptide chain-mediated translational stalling. Science 326, 1412–1415 (2009). 10.1126/science.1177662

48 Su, T. et al. Structural basis of l-tryptophan-dependent inhibition of release factor 2 by the TnaC arrest peptide. Nucleic Acids Res 49, 9539–9547 (2021). 10.1093/nar/gkab665

49 Rae, C. D., Gordiyenko, Y. & Ramakrishnan, V. How a circularized tmRNA moves through the ribosome. Science 363, 740–744 (2019). 10.1126/science.aav9370

50 Deeng, J. et al. Dynamic Behavior of Trigger Factor on the Ribosome. J Mol Biol 428, 3588–3602 (2016). 10.1016/j.jmb.2016.06.007

51 Jomaa, A., Boehringer, D., Leibundgut, M. & Ban, N. Structures of the E. coli translating ribosome with SRP and its receptor and with the translocon. Nat Commun 7, 10471 (2016). 10.1038/ncomms10471

52 Jomaa, A. et al. Structure of the quaternary complex between SRP, SR, and translocon bound to the translating ribosome. Nat Commun 8, 15470 (2017). 10.1038/ncomms15470

53 Wang, S. et al. The molecular mechanism of cotranslational membrane protein recognition and targeting by SecA. Nat Struct Mol Biol 26, 919–929 (2019). 10.1038/s41594-019-0297-8

54 Zivanov, J. et al. New tools for automated high-resolution cryo-EM structure determination in RELION-3. Elife 7 (2018). 10.7554/eLife.42166

55 Meng, E. C. et al. UCSF ChimeraX: Tools for structure building and analysis. Protein Sci 32, e4792 (2023). 10.1002/pro.4792

56 Liebschner, D. et al. Macromolecular structure determination using X-rays, neutrons and electrons: recent developments in Phenix. Acta Crystallogr D Struct Biol 75, 861–877 (2019). 10.1107/S2059798319011471

57 Casanal, A., Lohkamp, B. & Emsley, P. Current developments in Coot for macromolecular model building of Electron Cryo-microscopy and Crystallographic Data. Protein Sci 29, 1069–1078 (2020). 10.1002/pro.3791

58 McCoy, A. J., Fucini, P., Noegel, A. A. & Stewart, M. Structural basis for dimerization of the Dictyostelium gelation factor (ABP120) rod. Nat Struct Biol 6, 836–841 (1999). 10.1038/12296

59 Pall, S. et al. Heterogeneous parallelization and acceleration of molecular dynamics simulations in GROMACS. J Chem Phys 153, 134110 (2020). 10.1063/5.0018516

60 Huang, J. et al. CHARMM36m: an improved force field for folded and intrinsically disordered proteins. Nat Methods 14, 71–73 (2017). 10.1038/nmeth.4067

61 Noeske, J. et al. High-resolution structure of the Escherichia coli ribosome. Nat Struct Mol Biol 22, 336–341 (2015). 10.1038/nsmb.2994

62. Hess, B., Bekker, H., Berendsen, H. J. C. & Fraaije, J. G. E. M. LINCS: A linear constraint solver for molecular simulations. Journal of Computational Chemistry 18, 1463–1472 (1997). 10.1002/(SICI)1096-987X(199709)18:12<1463::AID-JCC4>3.0.CO;2-H

63 Darden, T., York, D. & Pedersen, L. Particle mesh Ewald: An N⋅log(N) method for Ewald sums in large systems. The Journal of Chemical Physics 98, 10089–10092 (1993). 10.1063/1.464397

64 Bussi, G., Donadio, D. & Parrinello, M. Canonical sampling through velocity rescaling. J Chem Phys 126, 014101 (2007). 10.1063/1.2408420

65 Berendsen, H. J. C., Postma, J. P. M., van Gunsteren, W. F., DiNola, A. & Haak, J. R. Molecular dynamics with coupling to an external bath. The Journal of Chemical Physics 81, 3684–3690 (1984). 10.1063/1.448118

66 Parrinello, M. & Rahman, A. Polymorphic transitions in single crystals: A new molecular dynamics method. Journal of Applied Physics 52, 7182–7190 (1981). 10.1063/1.328693

67 Hopkins, C. W., Le Grand, S., Walker, R. C. & Roitberg, A. E. Long-Time-Step Molecular Dynamics through Hydrogen Mass Repartitioning. J Chem Theory Comput 11, 1864–1874 (2015). 10.1021/ct5010406

68 Wong, V. & Case, D. A. Evaluating rotational diffusion from protein MD simulations. J Phys Chem B 112, 6013–6024 (2008). 10.1021/jp0761564

69 Wong, V., Case, D. A. & Szabo, A. Influence of the coupling of interdomain and overall motions on NMR relaxation. Proc Natl Acad Sci U S A 106, 11016–11021 (2009). 10.1073/pnas.0809994106

70 Lavalette, D., Amand, B. & Pochon, F. Rotational relaxation of 70S ribosomes by a depolarization method using triplet probes. Proc Natl Acad Sci U S A 74, 1407–1411 (1977). 10.1073/pnas.74.4.1407

71 Wlodarski, T. et al. Bayesian reweighting of biomolecular structural ensembles using heterogeneous cryo-EM maps with the cryoENsemble method. Sci Rep 14, 18149 (2024). 10.1038/s41598-024-68468-7

72 Briones, R., Blau, C., Kutzner, C., de Groot, B. L. & Aponte-Santamaría, C. GROma *ρ*s: A GROMACS-Based Toolset to Analyze Density Maps Derived from Molecular Dynamics Simulations. Biophysical Journal 116, 4–11 (2019). 10.1016/j.bpj.2018.11.3126

