## Supplementary information for "The ribosome directs nascent chains through two folding-dependent pathways"

**Supplementary Table S1. Cryo-EM data collection, refinement and validation statistics for FLN5+47 RNCs.**

| | FLN5+47 P-tRNA (EMDB-xxxx) (PDB xxxx) | FLN5+47 AP-tRNA (EMDB-xxxx) (PDB xxxx) | FLN5+47U (EMDB-xxxx) (PDB xxxx) |
|---|---|---|---|
| **Data collection and processing** | | | |
| Magnification | 81k | 81k | 130k |
| Voltage (kV) | 300 | 300 | 300 |
| Electron exposure (e–/Å$^2$) | 40 | 40 | 37 |
| Defocus range (μm) | 0.5-2.5 | 0.5-2.5 | 0.6-3 |
| Pixel size (Å) | 1.067/2 | 1.067/2 | 1.05 |
| Symmetry imposed | N/A | N/A | N/A |
| Initial particle images (no.) | 648,915 | 648,915 | 589,792 |
| Final particle images (no.) | 77,590 | 100,252 | 143,483 |
| Map resolution (Å)<br>FSC threshold 0.143 | 2.1 | 2.1 | 2.6 |
| **Refinement** | | | |
| Initial model used (PDB code) | 7zp8 | 7zp8 | 7zp8 |
| Model resolution (Å)<br>FSC threshold | 2.1/2.4/3.0<br>0/0.143/0.5 | 2.1/2.1/2.6<br>0/0.143/0.5 | 2.4/2.8/3.1<br>0/0.143/0.5 |
| Map sharpening *B* factor (Å$^2$) | -35 | -35 | -60 |
| Model composition<br>Non-hydrogen atoms | 32 chains<br>90,839 | 33 chains<br>92,476 | 32 chains<br>90,664 |

| | | | |
|---|---|---|---|
| Protein residues | 3,127 | 3,132 | 3,144 |
| Nucleotides | 3,096 | 3,171 | 3,085 |
| *B* factors (Å$^2$) | | | |
| Protein | 87 | 82 | 116 |
| Nucleotide | 96 | 97 | 127 |
| R.m.s. deviations | | | |
| Bond lengths (Å) | 0.009 | 0.011 | 0.010 |
| Bond angles (°) | 1.144 | 1.467 | 1.830 |
| Validation | | | |
| MolProbity score | 1.9 | 1.8 | 2.6 |
| Clashscore | 5.7 | 5.4 | 10.5 |
| Poor rotamers (%) | 2.2 | 1.3 | 7.5 |
| Ramachandran plot | | | |
| Favored (%) | 95.4 | 94.6 | 93.2 |
| Allowed (%) | 4.1 | 5.0 | 6.2 |
| Disallowed (%) | 0.5 | 0.4 | 0.6 |

**Supplementary Table S2. Cryo-EM data collection, refinement and validation statistics for FLN5+31 RNCs.**

| | FLN5+31 P-tRNA (EMDB-xxxx) (PDB xxxx) | FLN5+31 AP-tRNA (EMDB-xxxx) (PDB xxxx) | FLN5+31U (EMDB-xxxx) (PDB xxxx) |
|---|---|---|---|
| **Data collection and processing** | | | |
| Magnification | 105k | 105k | 165k |
| Voltage (kV) | 300 | 300 | 300 |
| Electron exposure (e–/Å$^2$) | 47 | 47 | |
| Defocus range (μm) | 0.3-2.1 | 0.3-2.1 | |
| Pixel size (Å) | 0.828/2 | 0.828/2 | 0.723/2 |
| Symmetry imposed | N/A | N/A | N/A |
| Initial particle images (no.) | 1,206,843 | 1,206,843 | 3,616,234 |
| Final particle images (no.) | 169,965 | 195,415 | 1,894,401 |
| Map resolution (Å)<br>FSC threshold 0.143 | 2.5 | 3.0 | 2.7 |
| **Refinement** | | | |
| Initial model used (PDB code) | 7zp8 | 7zp8 | 7zp8 |
| Model resolution (Å)<br>FSC threshold | 2.2/2.6/3.3<br>0/0.143/0.5 | 2.5/2.9/4.5<br>0/0.143/0.5 | <br>0/0.143/0.5 |
| Map sharpening *B* factor (Å$^2$) | | | |
| Model composition<br>Non-hydrogen atoms<br>Protein residues<br>Nucleotides | 32 chains<br>90,875<br>3,132<br>3,096 | 34 chains<br>92,670<br>3,132<br>3,180 | 32 chains |
| *B* factors (Å$^2$)<br>Protein<br>Nucleotide | <br>87<br>97 | <br>87<br>95 | |
| R.m.s. deviations<br>Bond lengths (Å)<br>Bond angles (°) | <br>0.009<br>1.164 | <br>0.008<br>1.045 | |
| Validation<br>MolProbity score<br>Clashscore<br>Poor rotamers (%) | <br>1.7<br>5.8<br>1.3 | <br>1.9<br>6.3<br>1.8 | |
| Ramachandran plot<br>Favored (%)<br>Allowed (%)<br>Disallowed (%) | <br>95.2<br>4.6<br>0.2 | <br>94.9<br>4.8<br>0.3 | |

**SupplementaryTable S3. Cryo-EM data collection, refinement and validation statistics for FLN5+34 RNC.**

| | FLN5+34 AP-tRNA<br>(EMDB-xxxx)<br>(PDB xxxx) |
|---|---|
| **Data collection and processing** | |
| Magnification | 81k |
| Voltage (kV) | 300 |
| Electron exposure (e–/Å$^2$) | 43 |
| Defocus range (μm) | 1.2-2.4 |
| Pixel size (Å) | 1.072/2 |
| Symmetry imposed | N/A |
| Initial particle images (no.) | |
| Final particle images (no.) | 281,242 |
| Map resolution (Å)<br>FSC threshold | 2.2 |
| **Refinement** | |
| Initial model used (PDB code) | 7zp8 |
| Model resolution (Å)<br>FSC threshold | 2.1/2.3/3.1<br>0/0.143/0.5 |
| Map sharpening *B* factor (Å$^2$) | -34 |
| Model composition<br>Non-hydrogen atoms<br>Protein residues<br>Ligands | <br>92,691<br>3,132<br>3,181 |
| *B* factors (Å$^2$)<br>Protein<br>Ligand | <br>87<br>96 |
| R.m.s. deviations<br>Bond lengths (Å)<br>Bond angles (°) | <br>0.011<br>1.256 |
| Validation<br>MolProbity score<br>Clashscore<br>Poor rotamers (%) | <br>1.8<br>6.4<br>1.4 |
| Ramachandran plot<br>Favored (%)<br>Allowed (%)<br>Disallowed (%) | <br>95.2<br>4.6<br>0.2 |

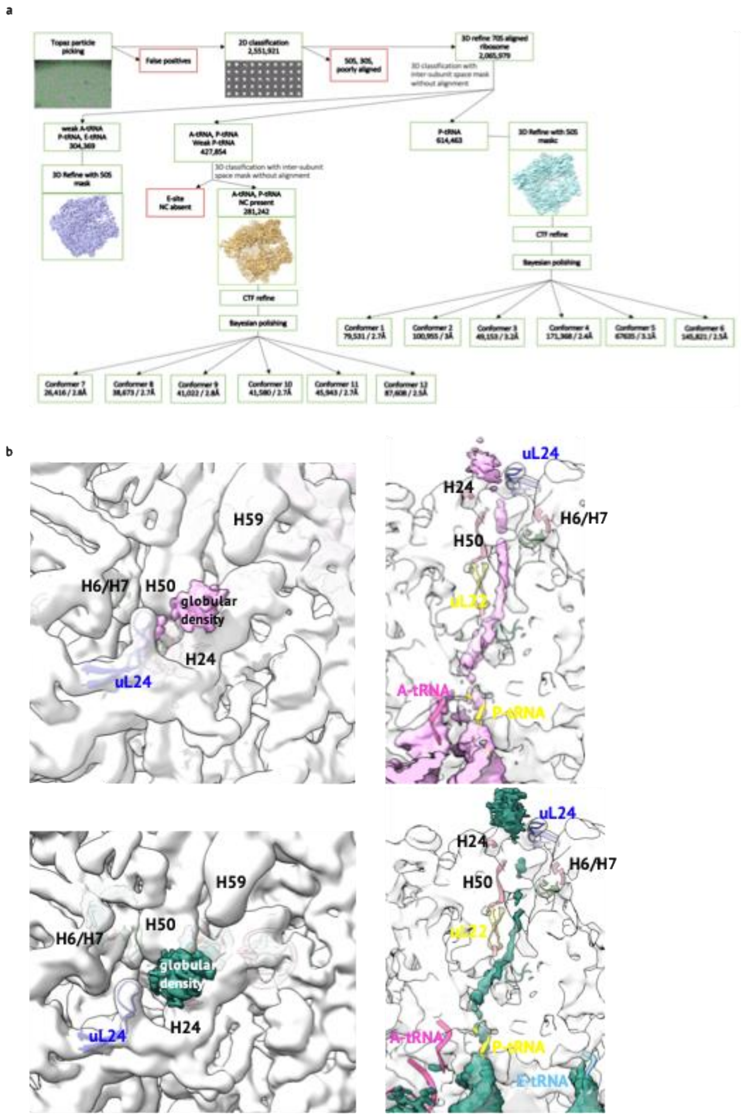

**Supplementary Fig. 1**. **a.** In silico particle sorting for FLN5+34 RNC. From 11,502 micrographs, approximately 1.8 million particles were picked and subjected to further processing. Discarded particles are depicted in red boxes while green boxes show particles that were subjected to further processing. **b.** Reconstruction of AP-tRNA (top) and APE-tRNA (bottom) FLN5+34 before tunnel classification. (Left) Density for the FLN5 globular domain (pink and green surface representation for AP-tRNA and APE-tRNA RNCs, respectively) is observed beyond the tunnel near uL24. (Right) Transverse section of the reconstructions shows the NC density in the tunnel up to uL23, with discontinuous density in the vestibule, reappearing beyond the tunnel. The reconstructions are shown in white surface representation for the 50S ribosomal subunit.

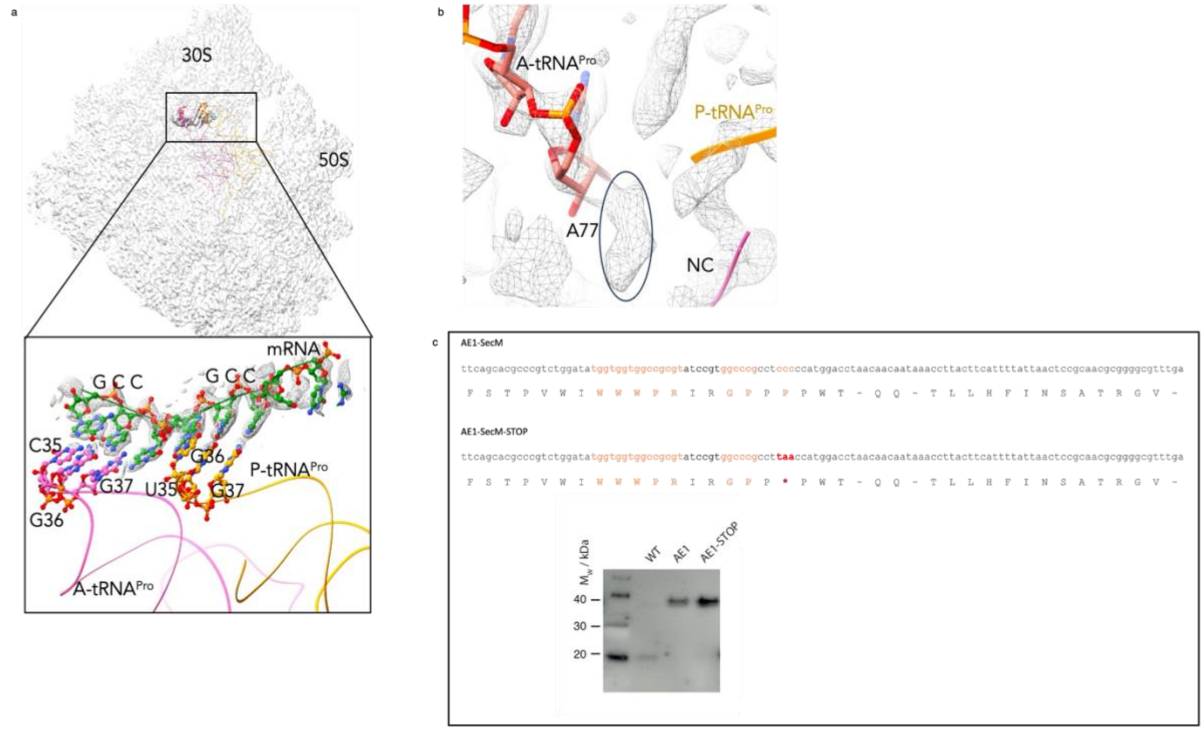

**Supplementary Fig. 2**. **a**. Pre-translocation states of arrest-enhanced SecM RNCs. Overall view of the AP-tRNA RNC reconstruction at 2.2Å (white surface), with A-site tRNA$^{pro}$ (magenta) and P-site tRNA$^{pro}$ (yellow models). The boxed region corresponds to the decoding region. Zoomed-in view of the decoding region with the map (grey mesh) shown at 3 σ level. The de-novo-built mRNA model is shown in green and the anticodon bases of A- and P-site tRNA$^{pro}$ are shown in magenta and yellow, respectively. **b**. Cryo-EM density at 2.5 σ level of AP-tRNA RNC, showing the CCA-end of A- tRNA$^{pro}$ (depicted in pink atoms) attached to a density which is attributed to the proline amino acid. Cartoon representation of the CCA end with the first two amino acids of the NC shown in cartoon (magenta), which show continuous density from the CCA end to the NC, suggesting the NC is attached to the P- tRNA$^{pro}$. **c**. DNA and protein sequence of the incorporated enhanced SecM sequence (AE1) and AE1 with a stop codon after the stalling motif. (Bottom) Anti-histidine western blot analysis of cell extracts following expression of FLN5+31 RNC. The NC is stalled using SecM (WT), AE1, or AE1 with stop codon (AE1-STOP). The amount of release species (detected at around 20kDa) is higher for WT than the other two constructs, suggesting that the WT SecM is a weaker translation arrest peptide.

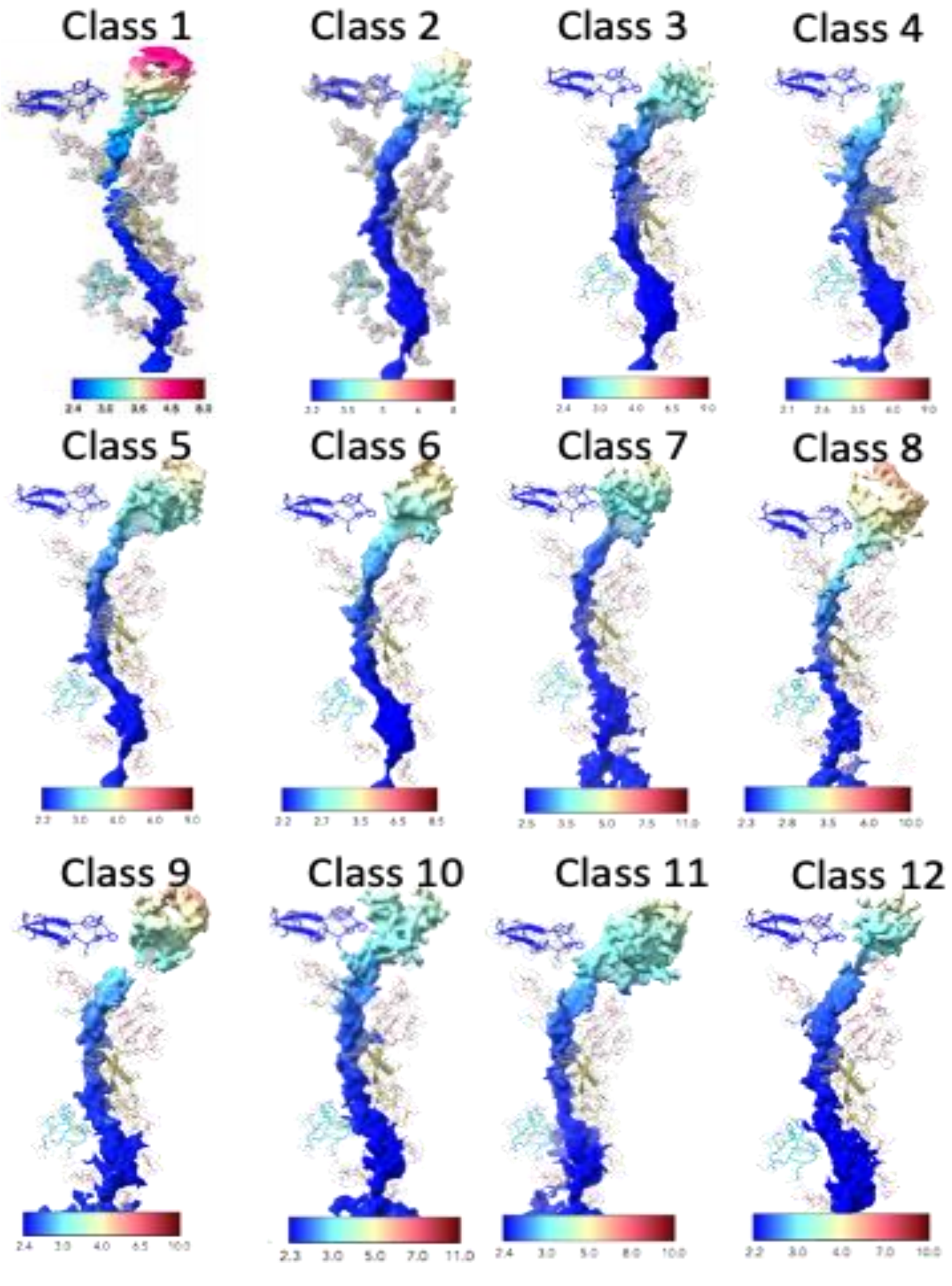

**Supplementary Fig. 3**. Local resolution for FLN5+34 RNCs after tunnel classification. The panels show difference maps for tRNA(s) and the nascent chain, highlighting variations in the resolutionacross different classes. Ribosomal landmarks are shown and labeled as follows: uL4, uL22, uL23 and uL24 loops are shown in cyan, yellow, light green and blue, respectively. rRNA bases A64, A91, C490, A751, C1320-G1324, A2062, U2585 and U2609 are shown in pink (as shown in Fig 2b).

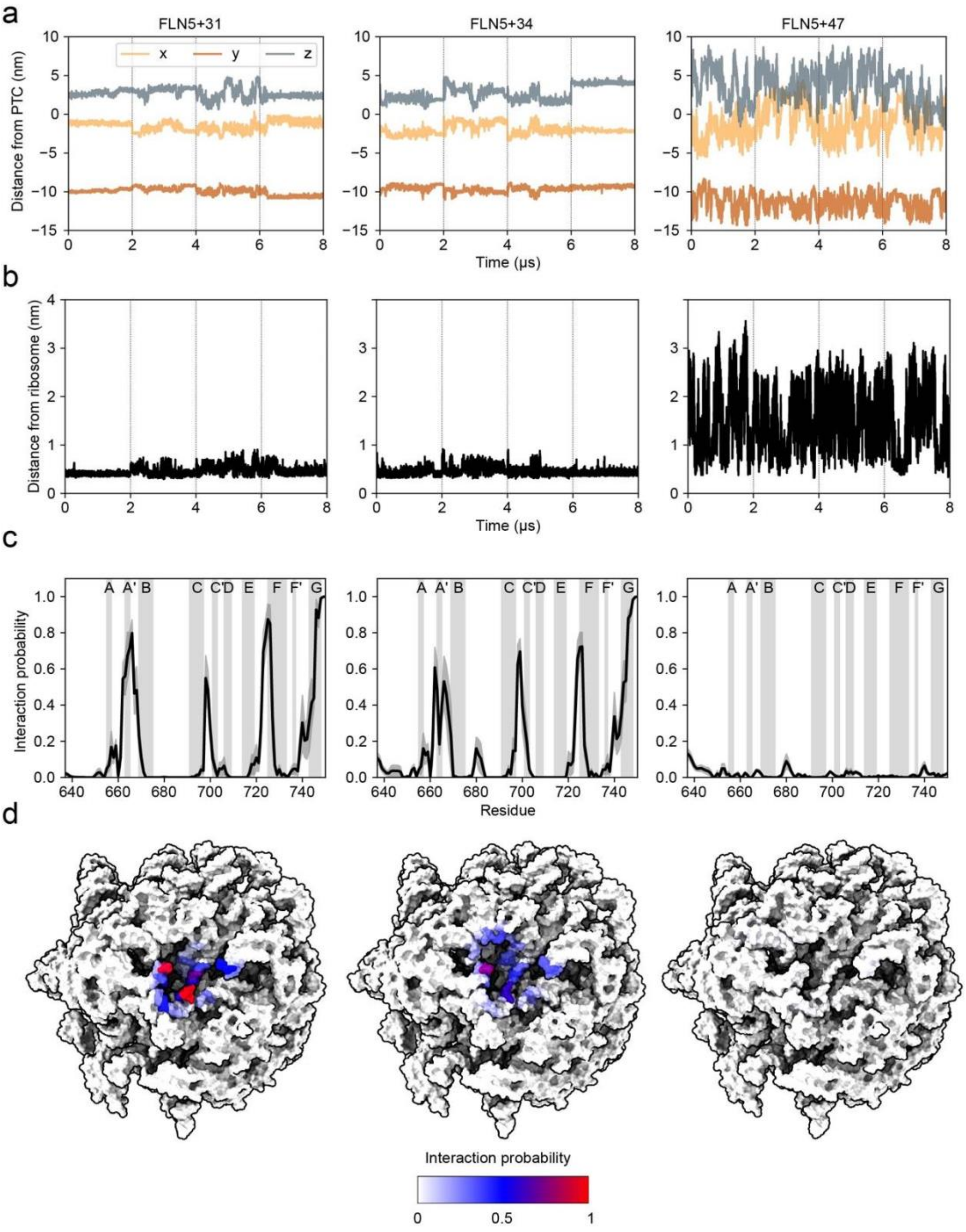

**Supplementary Fig. 4**. Dynamics and ribosome interactions of folded FLN5 observed in all-atom (C36m+W) MD simulations on the ribosome at three linker lengths. **(a)** Components of the distance between the centre of mass of FLN5 and the PTC (C-terminal proline, Cα position) calculated from four independent simulations of 2μs (individual trajectories separated by the vertical lines). **(b)** Minimum distance between FLN5 (Cα atoms) and the ribosome (Cα, P, N3, C4' atoms) calculated from four independent 2μs simulations. **(c)** Ribosome interaction probability plotted along the sequence of FLN5 for each RNC length (mean ± SEM from 4x independent 2μs simulations). β-strand locations are annotated on the plots. **(d)** FLN5 interaction probabilities mapped onto the ribosome surface, calculated as an average across all four 2μs simulations.

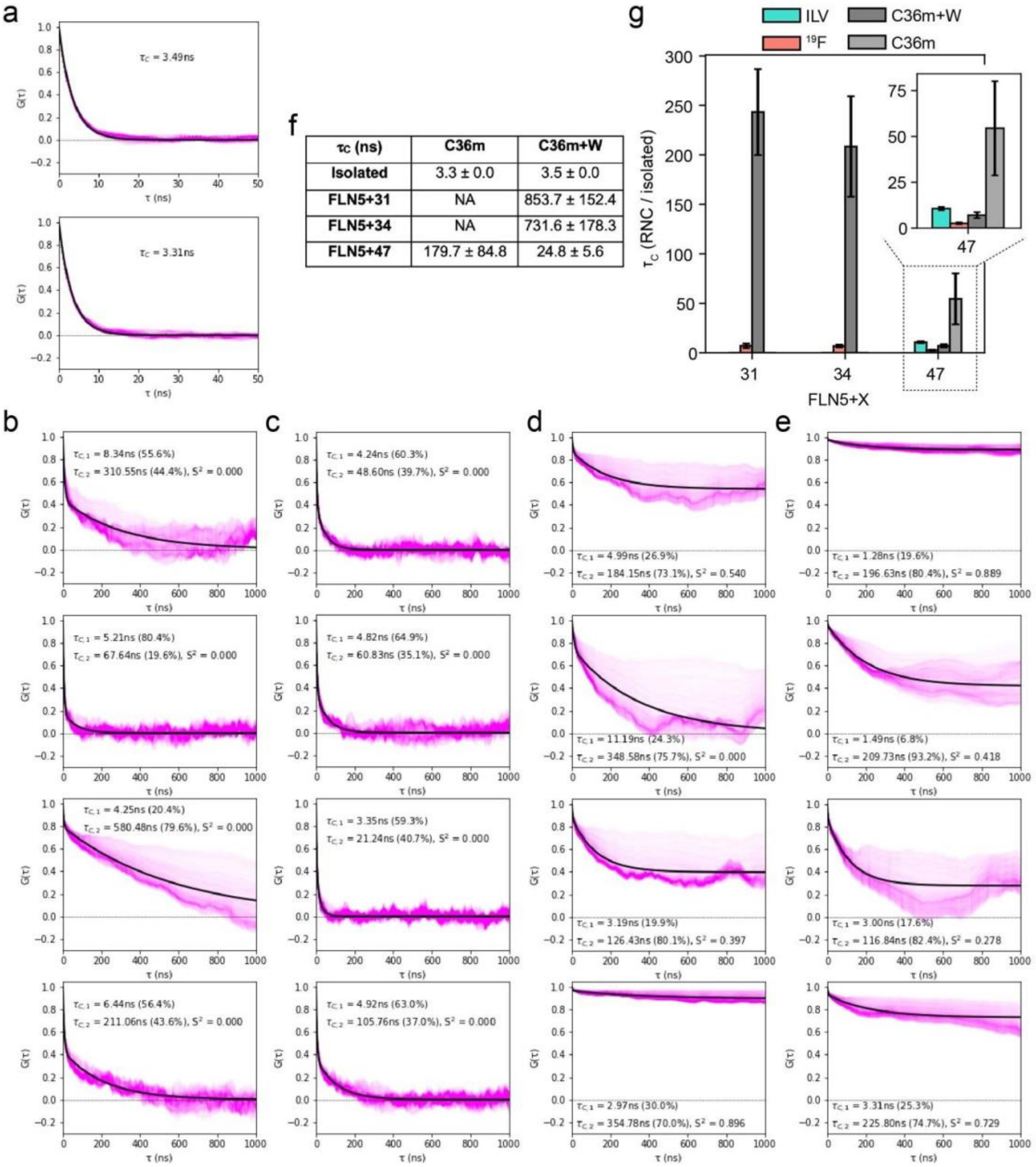

| τc (ns) | C36m | C36m+W |
|---|---|---|
| **Isolated** | 3.3 ± 0.0 | 3.5 ± 0.0 |
| **FLN5+31** | NA | 853.7 ± 152.4 |
| **FLN5+34** | NA | 731.6 ± 178.3 |
| **FLN5+47** | 179.7 ± 84.8 | 24.8 ± 5.6 |

**Supplementary Fig. 5**. Comparison of the FLN5 rotational correlation times ($\tau_C$) predicted by all-atom MD simulations with NMR-measured values (see Methods). **(a)** Exemplar rotational correlation function plots (global fit, black, to calculated correlation functions of 1,000 uniformly distributed vectors in the unit sphere, magenta) of isolated FLN5 simulated with C36m+W (top) and C36m (bottom). **(b-e)** Rotational correlation function plots (fit and calculated from MD) of FLN5 RNCs: **(b)** FLN5+47, C36m; **(c)** FLN5+47, C36m+W; **(d)** FLN5+34, C36m+W; **(e)** FLN5+31, C36m+W. MD data were fit to two correlation times and an order parameter (shown on the plot, see Methods). **(f)** FLN5 rotational correlation times calculated from MD simulations (mean ± SEM from four independent 2μs simulations). **(g)** Bar plot showing MD- and NMR-derived rotational correlation times on the ribosome, normalised by their respective values for the isolated protein.

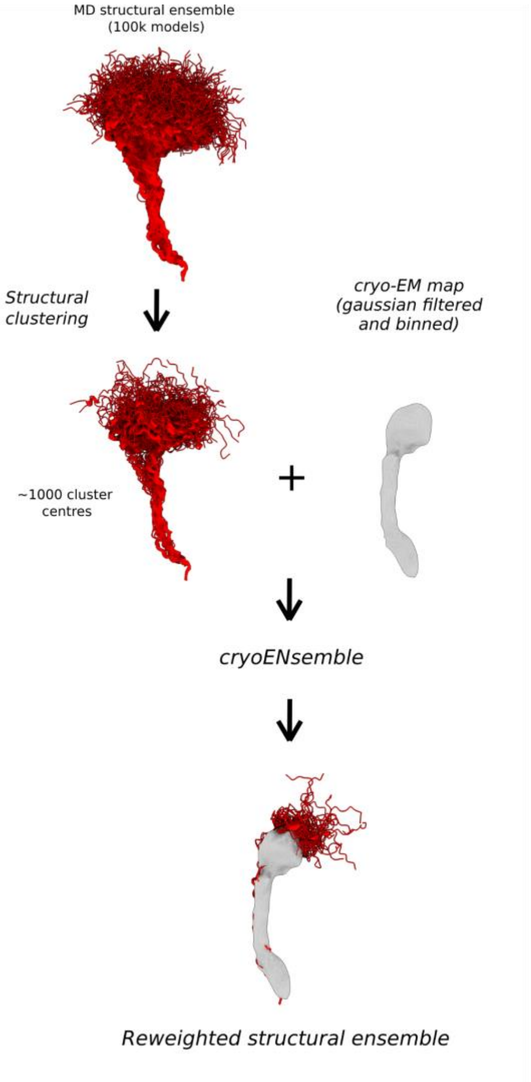

**Supplementary Fig. 6.** *Schematic of the cryoENsemble methodology used in the study. Structural clustering is performed in GROMACS using the gromos algorithm, yielding 1000 cluster centres. These centres are then reweighted in cryoENsemble using a Gaussian-filtered (with 3σ) and binned (2x) cryo-EM map. CryoENsemble identifies a structural sub-ensemble with new weights that improve the agreement with the corresponding cryo-EM map.*

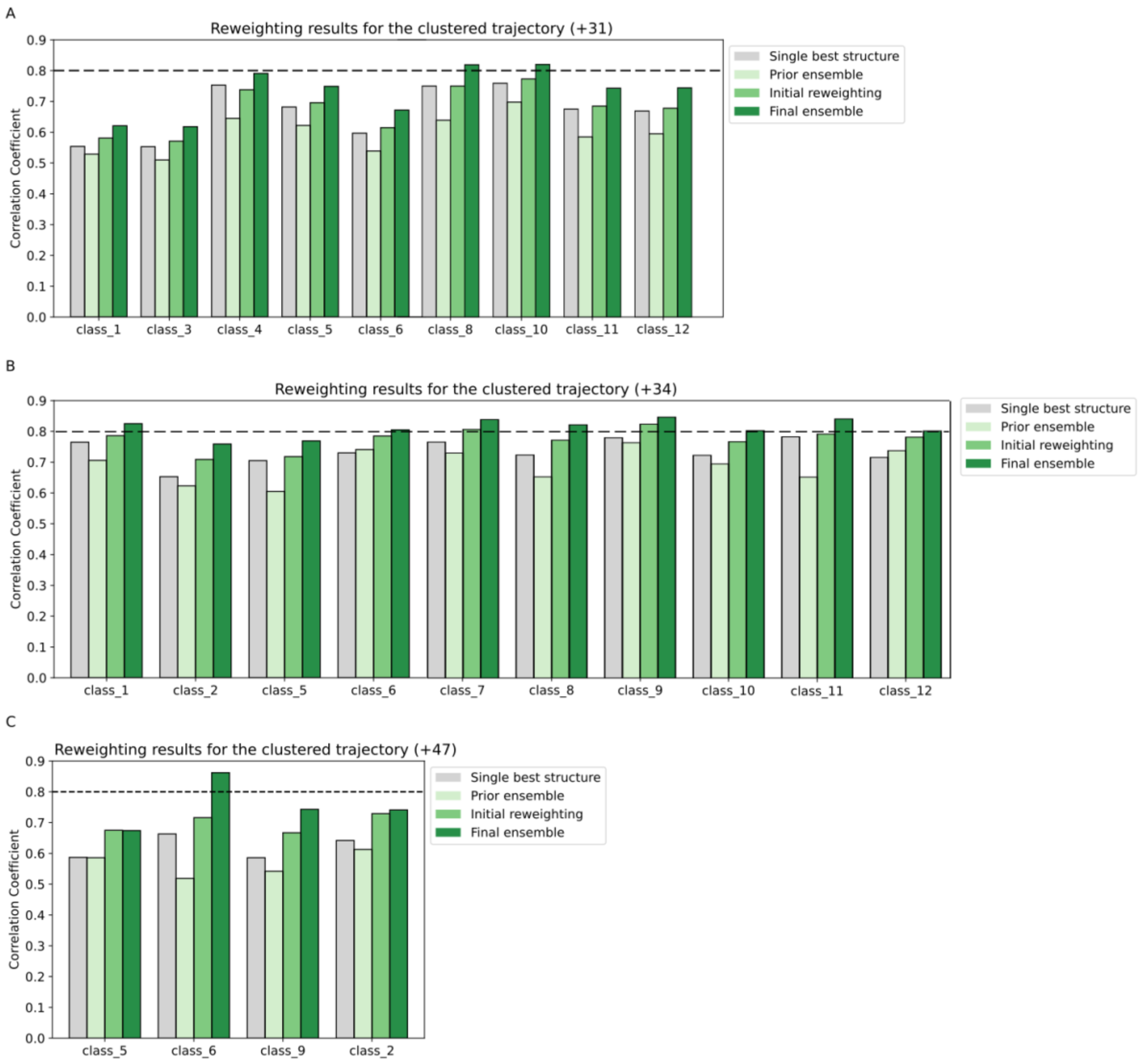

***Supplementary Fig. 7.*** *Cross-correlation coefficients obtained for the single best structure, prior ensemble, and after the first and last rounds of MD ensemble reweighting for (A) FLN5+31, (B) FLN5+34 and (C) FLN5+47.*

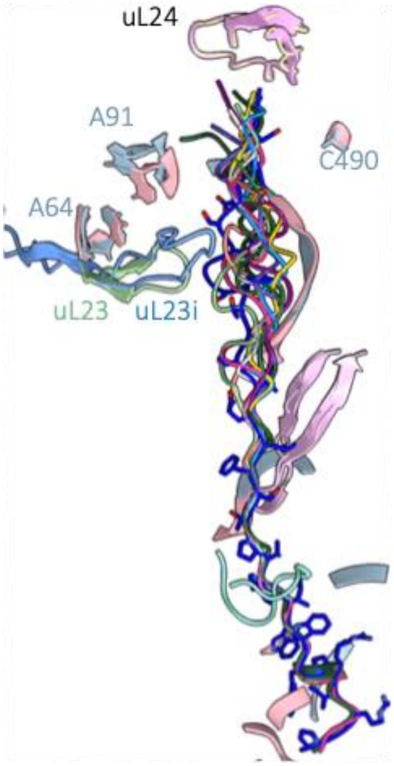

**Supplementary Fig. 8.** Comparison between the FLN5+34 RNCs and the RNC of the CRISPR-modified ribosome with FLN5 at L=34 (uL23 insertion) from Ahn et al. (in blue). Ribosomal landmarks within the exit tunnel are shown: uL22 and uL24 proteins are depicted in pink and yellow for FLN5+34 and 23i_FLN5+34, respectively. The 23S rRNA bases A64-A90 (H6-H7 rRNA helices), C490 (H24 rRNA helix) and A1321 (H50 rRNA helix) are shown in pink atoms. The modified uL23 loop is depicted in light blue and the wild-type loop in light green.

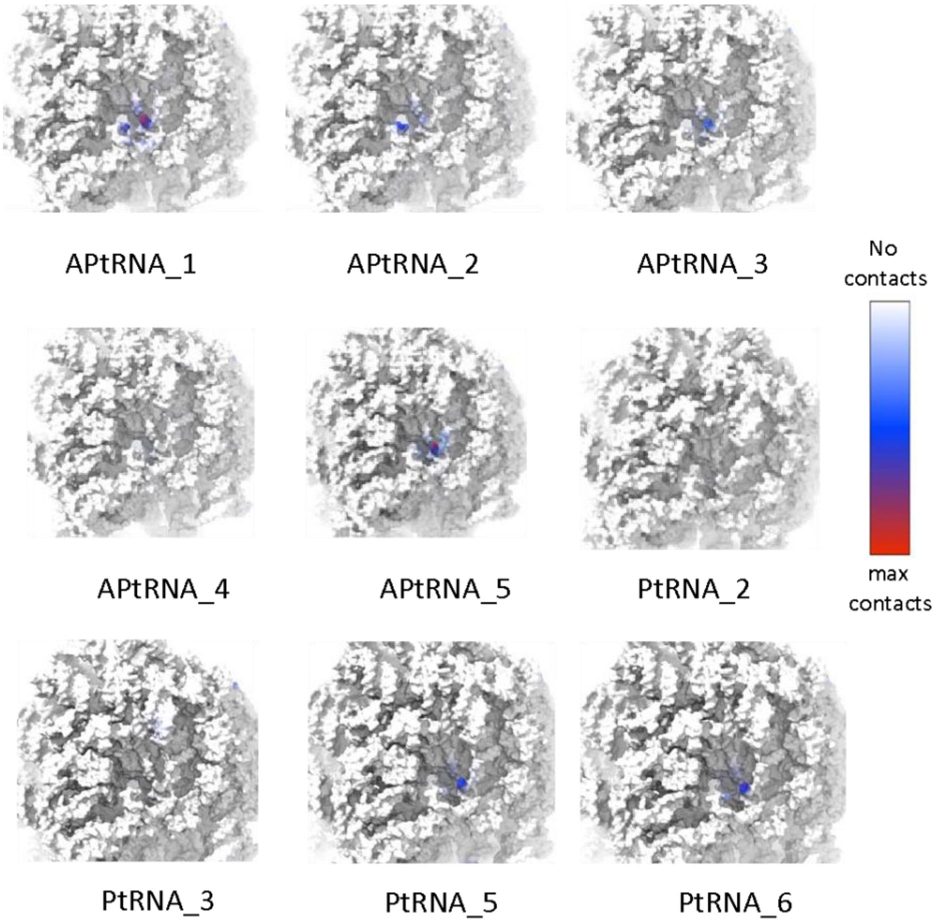

**Supplementary Fig. 9.** Calculated contacts between the ribosomal vestibule and the NC density for FLN5+34 RNCs. White corresponds to zero NC density voxel observed within 10 Å from the vestibule, while red represents the maximum number of voxels observed near the ribosomal surface. Note that these contacts do not correspond to NC density outside the tunnel as they are located below the uL24 loop which marks the end of the exit tunnel.

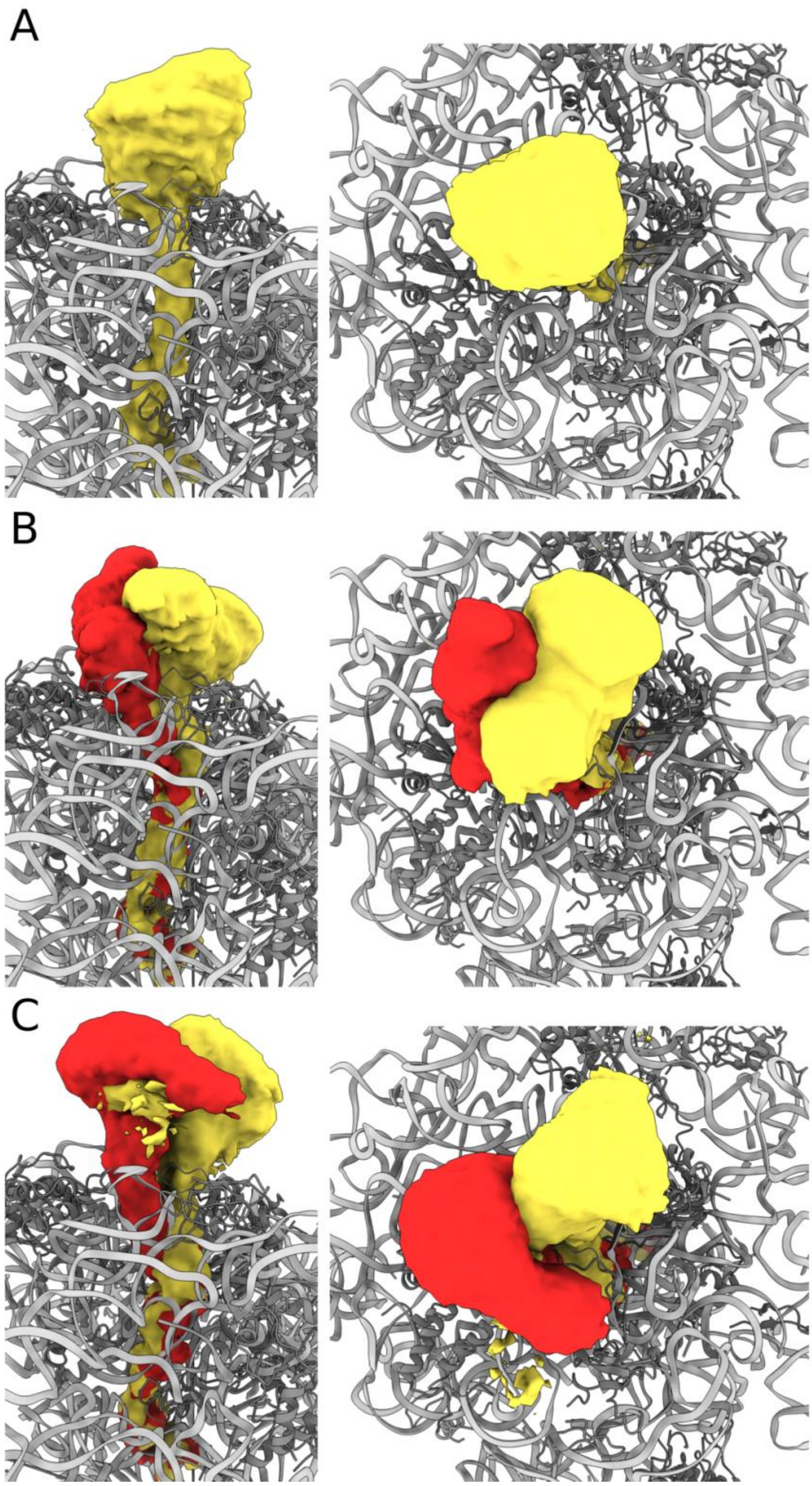

**Supplementary Fig. 10**. Density maps calculated using GROmaps[72] based on subensembles from MD simulations: A) FLN5+31 RNC, B) FLN5+34 RNC, C) FLN5+47 RNC. Red indicates the density corresponding to structures that use the path H6/H7, while yellow represents the H24/H50 pathway.

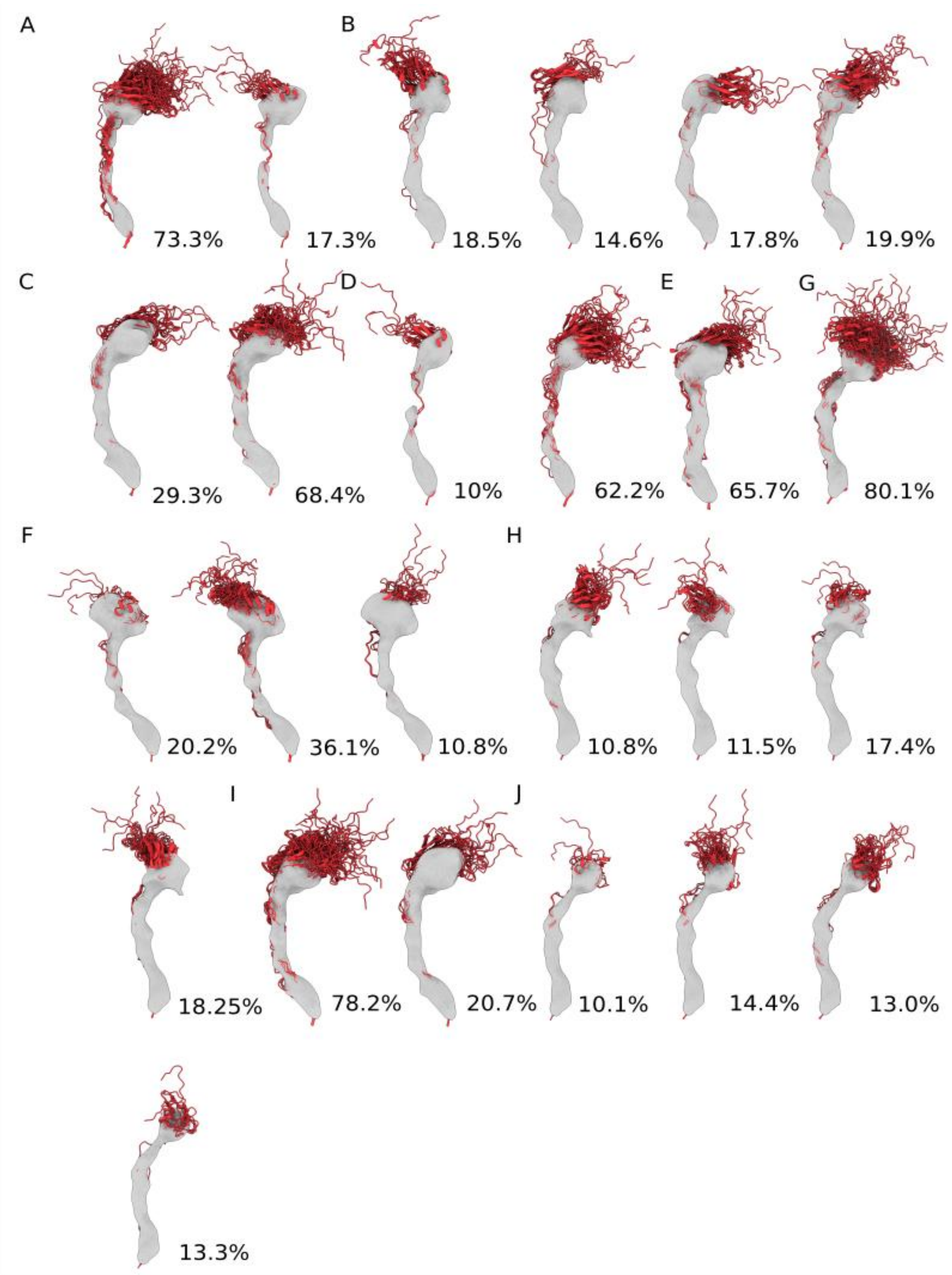

**Supplementary Fig. 11.** *Structural ensembles corresponding to the main states (with a population > 10%) generated upon reweighting the +34 RNC MD ensembles. Ensembles A through J correspond to the classes_1, 2, 5, 6, 7, 8, 9, 10, 11 and 12. The corresponding NC maps, Gaussian-filtered at the 3σ level, are visualised along with the population of each state.*

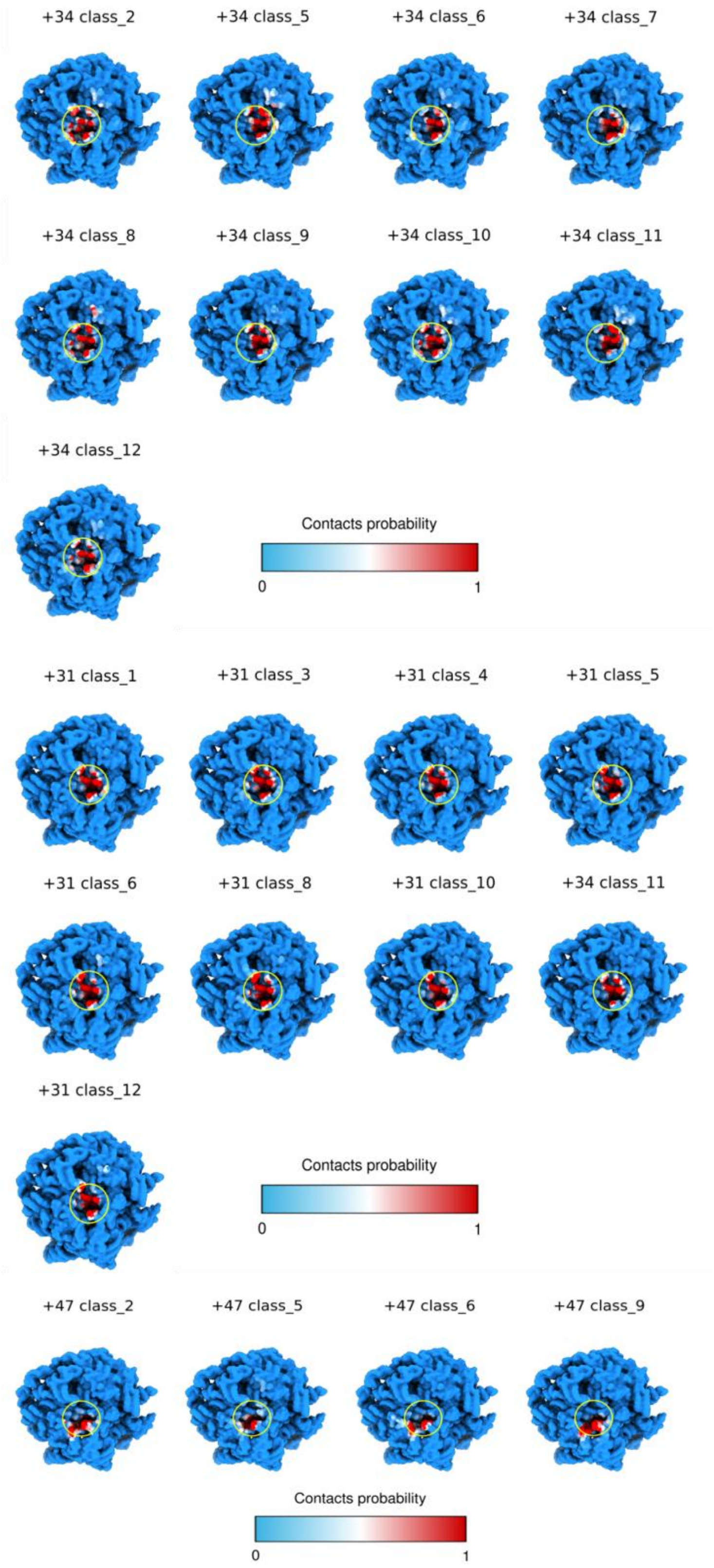

**Supplementary Fig. 12.** *Ribosome surface colored based on NC contact probability for reweighted MD ensemble from FLN5+34 (A), FLN5+31 (B) and FLN5+47 (C) FLN RNCs.*

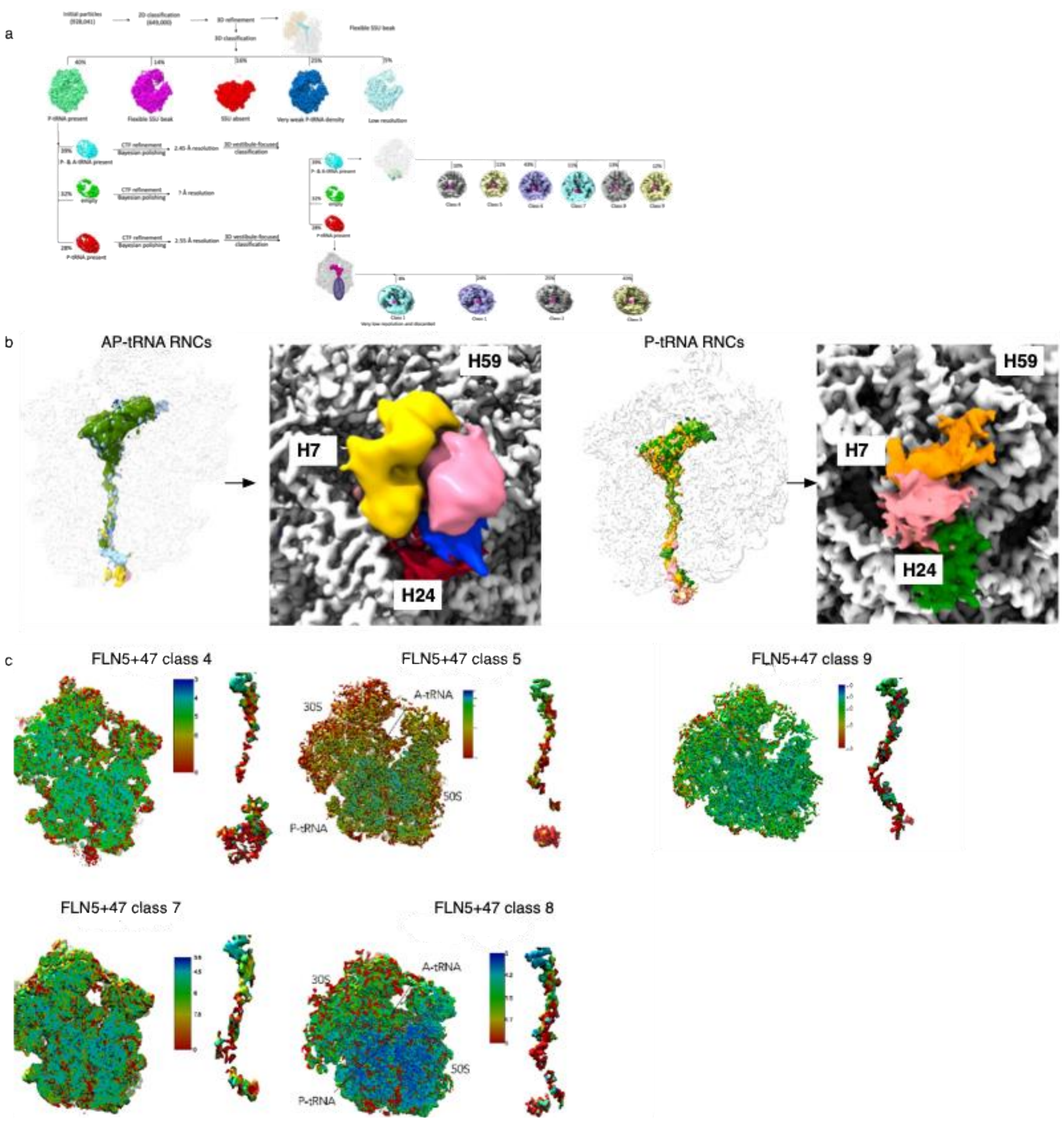

**Supplementary Fig. 13**. In silico particle sorting for FLN5+47 RNC. From 3,500 micrographs, over one million particles were picked and subjected to further processing. Vestibule 3D classification is shown on the right. b. Transverse and aerial view of FLN5+47 RNCs; ribosome density (grey surface), and the NC maps are coloured by class. c. Local resolution density maps for FLN5+47 RNCs after tunnel classification. The panels show 70S and representative difference maps for the FLN5+47 classes. Note the bar varies between different classes.

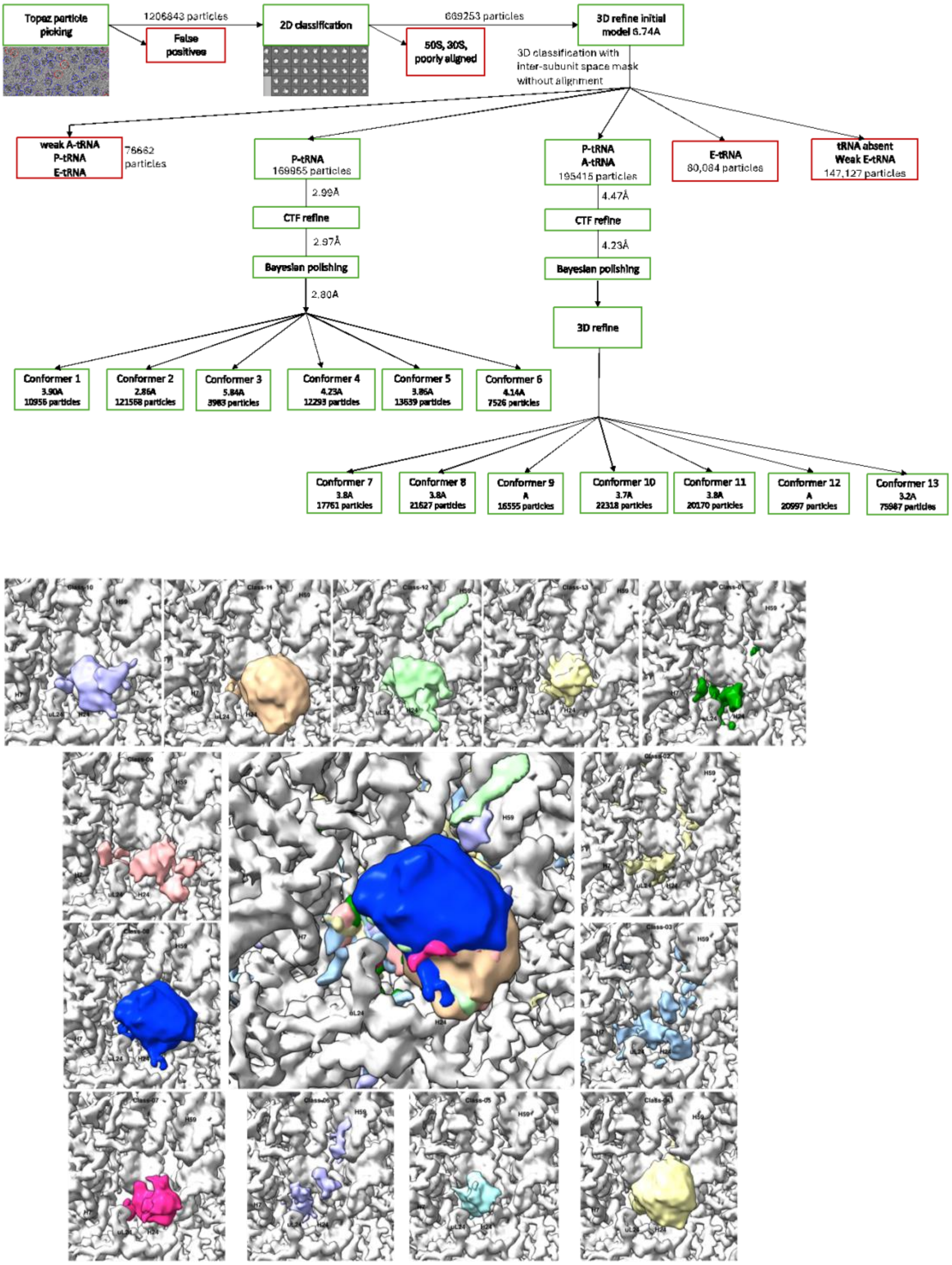

**Supplementary Fig. 14**. a) In silico particle sorting for FLN5+31 RNC. From 12,000 micrographs, approximately 1.2 million particles were picked and subjected to further processing. Discarded particles are depicted in red boxes, while green boxes show particles that were subjected to further processing. b) Aerial view of FLN5+31 RNCs, showing ribosome density (grey surface), and NC maps coloured by class.(Inset) A series of classes and the locations of globular density outside the ribosome.

a

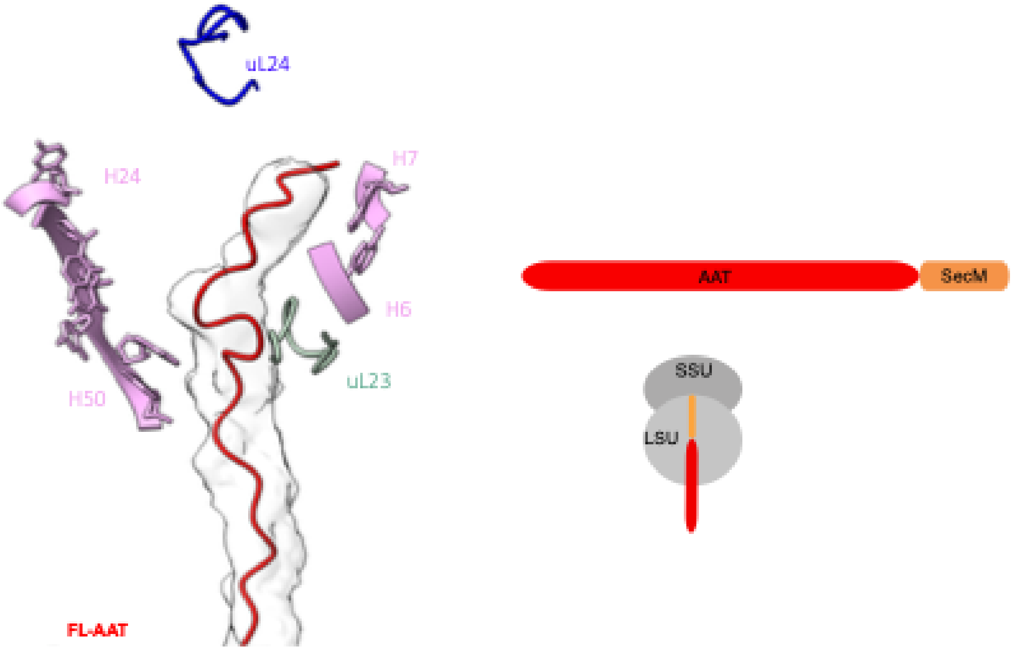

**Supplementary Fig. 15. Alpha-1 antitrypsin (AAT) RNC system as a model to study the folding pathway of expanded structures.** (Left) *The full-length AAT NC (model in red cartoon, density in white surface representations) is attached to arrest-enhanced SecM within the ribosomal exit tunnel. Key ribosomal landmarks are shown: uL23 (light green) and uL24 (blue), 23S rRNA bases A64-A90 (H6-H7 rRNA helices), C490 (H24 rRNA helix) and A1321 (H50 rRNA helix) in pink atoms. (Right) Schematic of the AAT construct used for this study.*

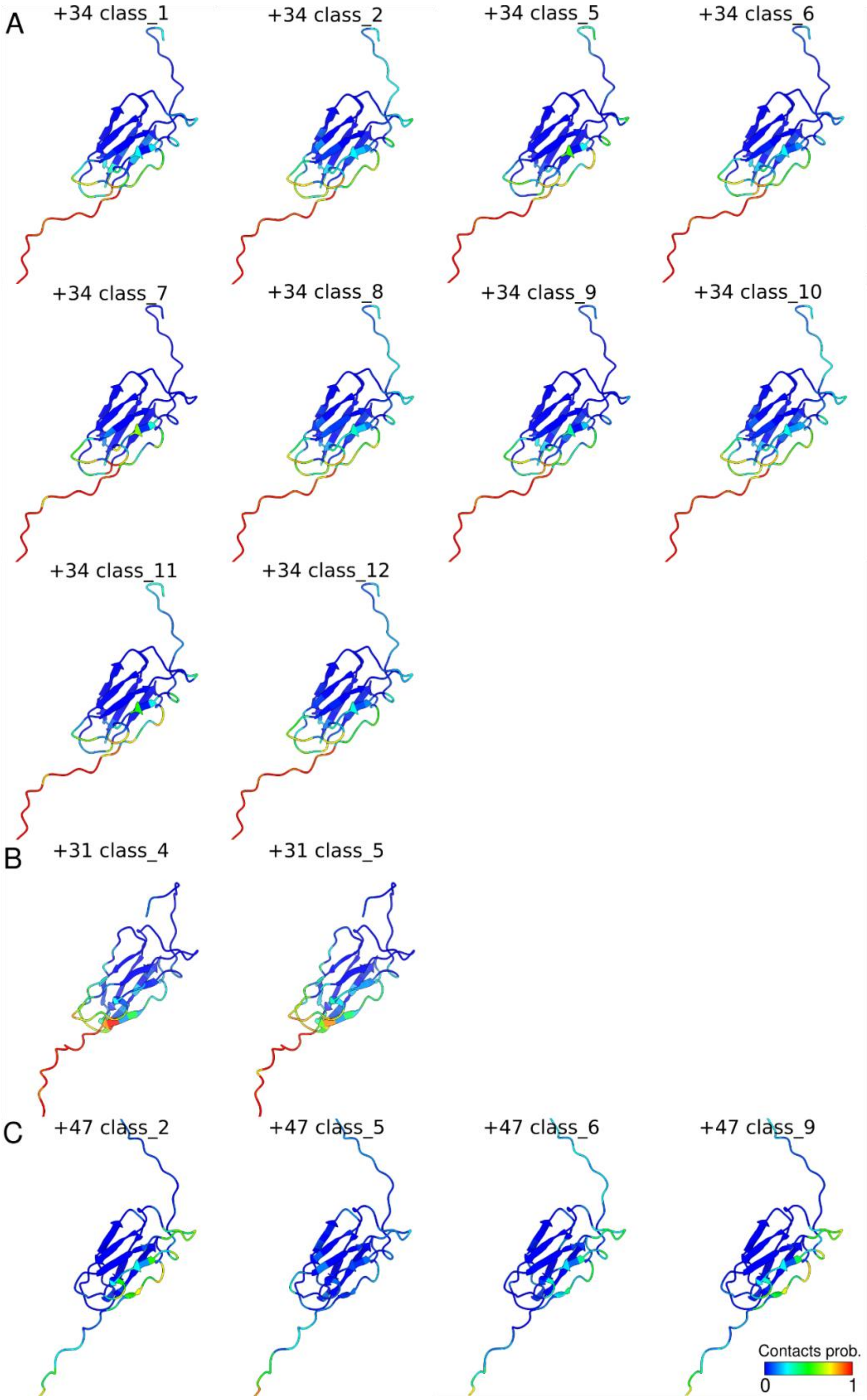

**Supplementary Fig. 16.** *Structures of the FLN5 NC colored according to the probability of contact with the ribosome surface, based on the reweighted MD ensembles for (A) FLN5+34, (B) FLN5+31 and (C) FLN5+47.*

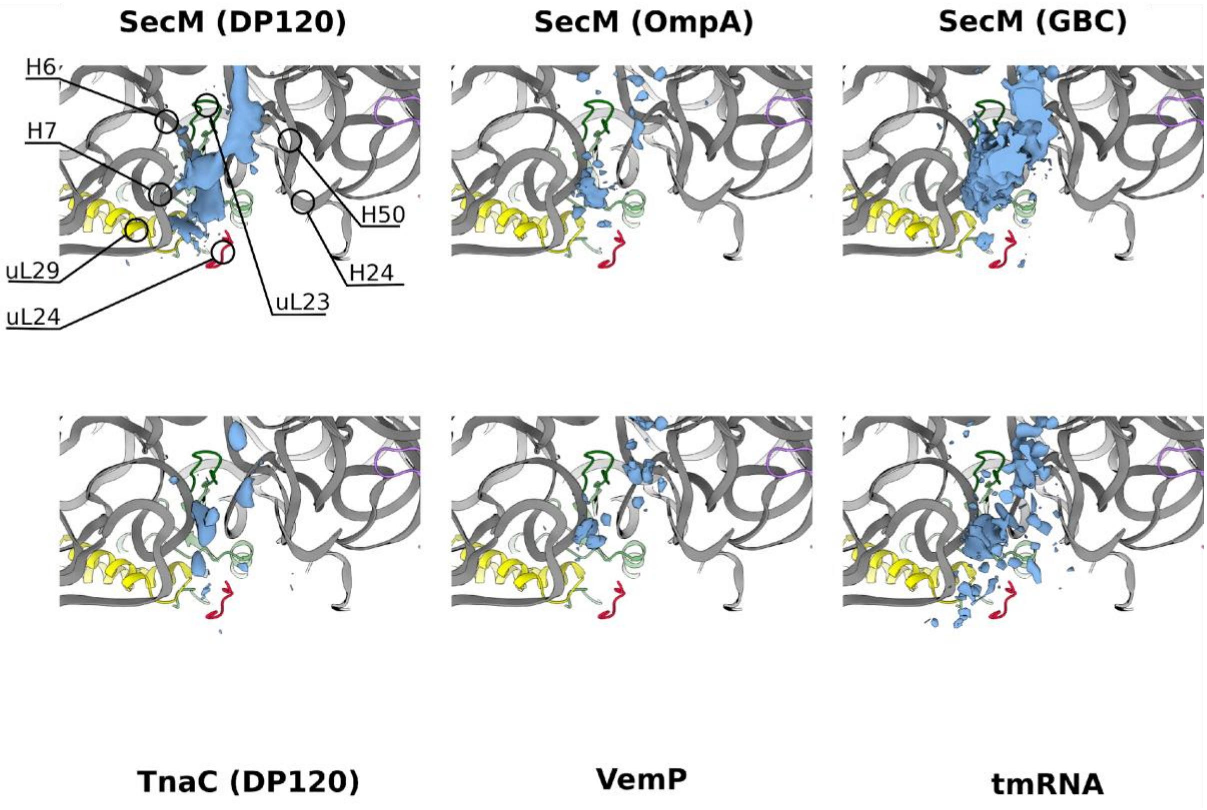

**Supplementary Fig. 17.** NC cryo-EM densities (in blue) obtained from various published *E. coli* stalled systems (see Suppl. Table 3), aligned with a single ribosome structure (PDB ID: 3J9Z) to enable comparison. NC cryo-EM densities are depicted using contour levels as recommended by the respective authors.

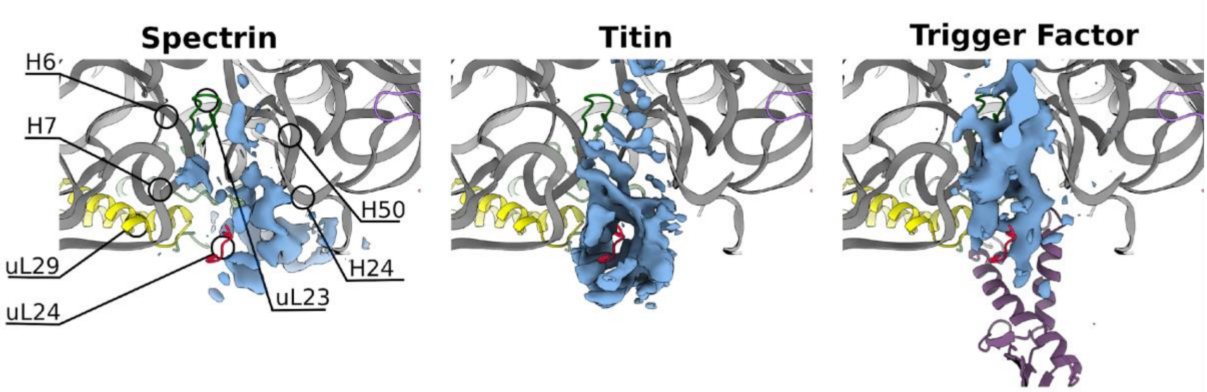

**Supplementary Fig. 18.** NC cryo-EM densities (in blue) obtained from two folded RNCs, spectrin[32] and titin[38], and from trigger factor (TF) bound to the ribosome[13] (RBD domain of the TF is in purple). All structures are aligned with a single ribosome structure (PDB ID: 3J9Z) for comparison. NC cryo-EM densities are depicted using contour levels as recommended by the respective authors.

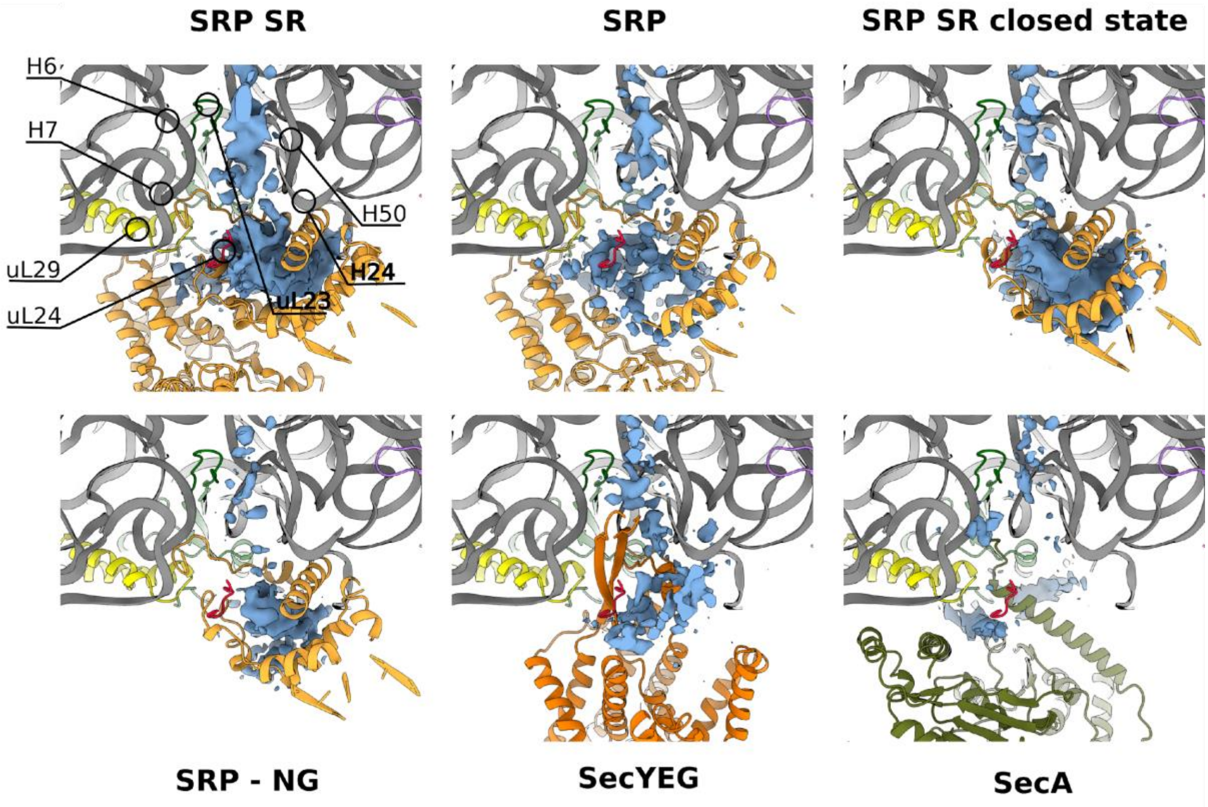

**Supplementary Fig. 19.** NC cryo-EM densities (in blue) obtained from several ribosome-associated factors involved in translocation (see Suppl. Table 3 for details and references), aligned with a single ribosome structure (PDB ID: 3J9Z) for comparison. NC cryo-EM densities are depicted using contour levels as recommended by the authors of these structures.

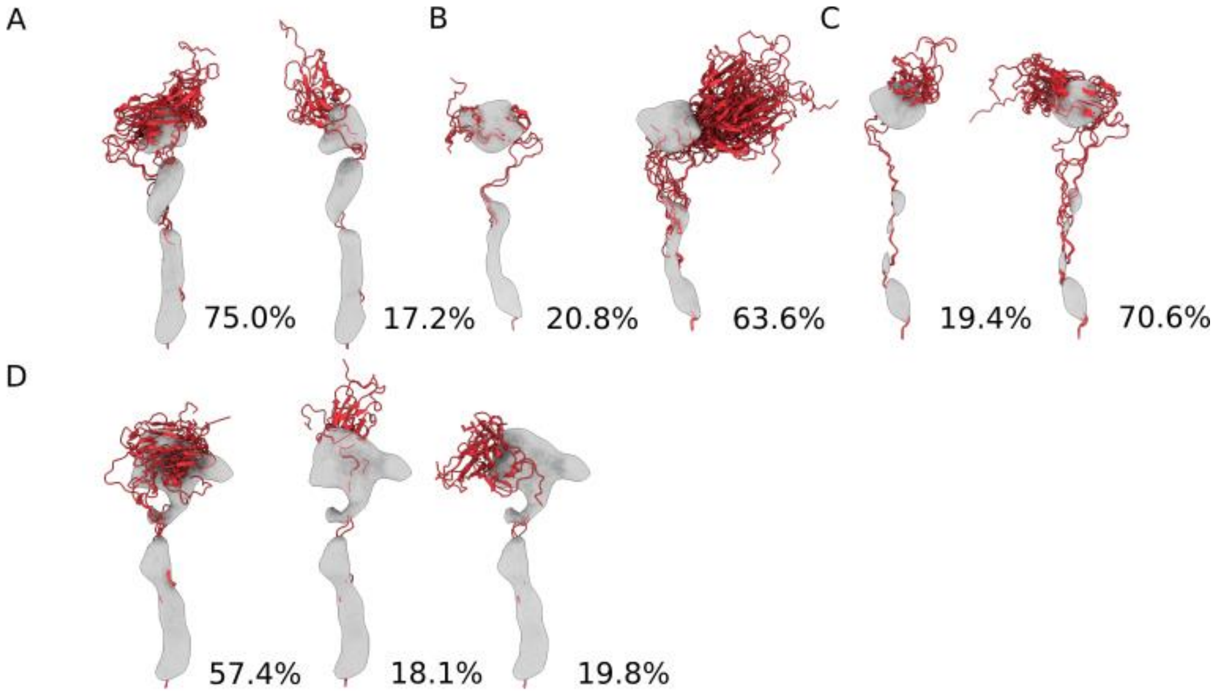

**Supplementary Fig. 20.** Structural ensembles corresponding to the main states (with a population > 10%) generated upon reweighting the +47 RNCs MD ensemble. Ensembles A through D correspond to the classes_2, 5, 6 and 9. The corresponding NC maps, Gaussian-filtered at the 3σ level, are visualised alongside the population of each state.

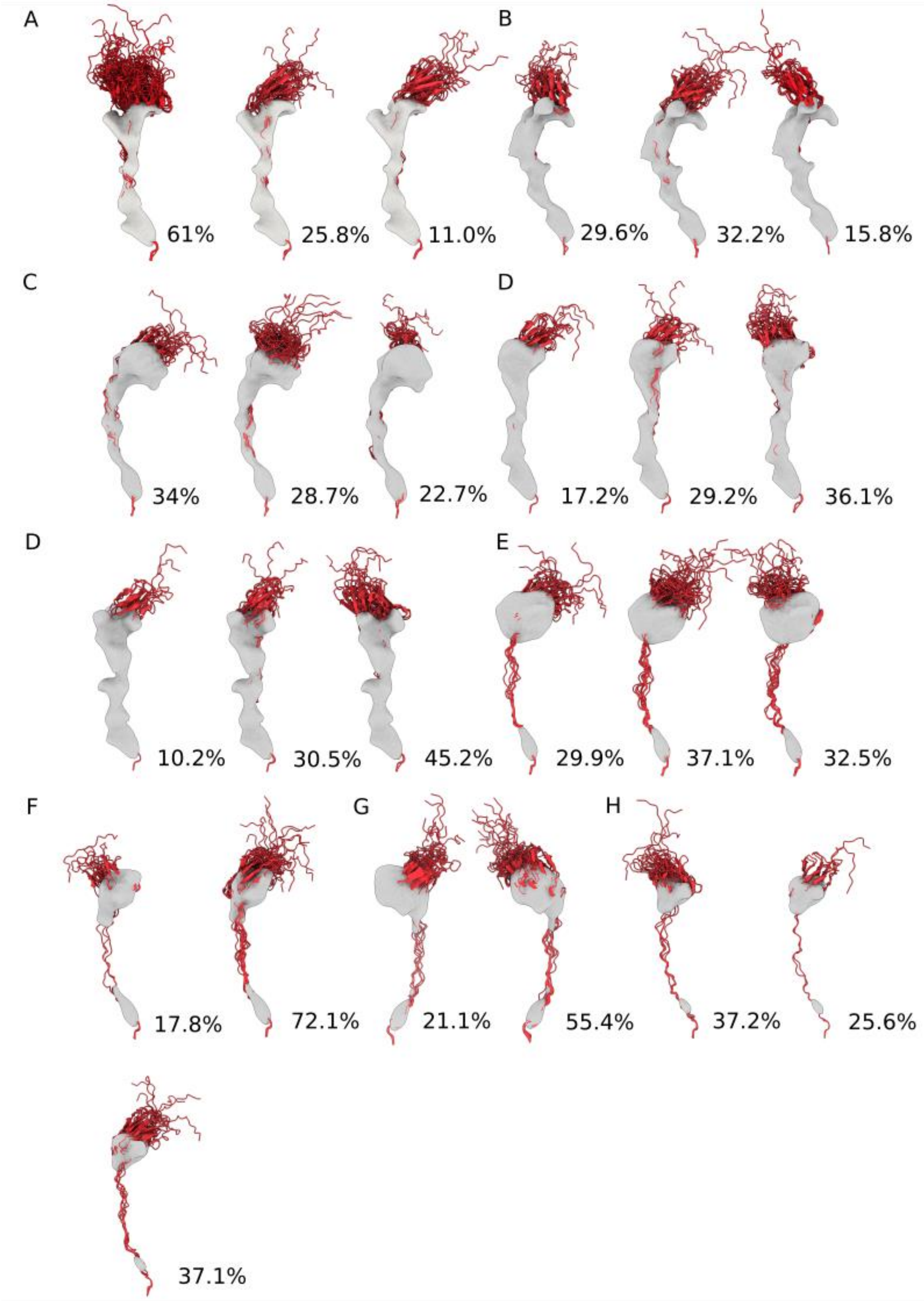

**Supplementary Fig. 21.** *Structural ensembles corresponding to the main states (with a population > 10%) generated upon reweighting the +31 RNCs MD ensemble. Ensembles A through H correspond to the classes_1, 3, 4, 5, 6, 8, 10, 11, and 12. The corresponding NC maps, Gaussian-filtered at the 3σ level, are visualised alongside the population of each state.*
